# On the dynamics of reproductive values and phenotypic traits in class-structured populations

**DOI:** 10.1101/155879

**Authors:** Sébastien Lion

## Abstract

In natural populations, individuals of a given genotype may belong to different classes. Such classes can for instance represent different age groups, developmental stages, or habitats. Class structure has important evolutionary consequences because the fitness of individuals with the same genetic background may vary depending on their class. As a result, demographic transitions between classes can cause fluctuations that need to be removed when estimating selection on a trait. Intrinsic differences between classes are classically taken into account by weighting individuals by class-specific reproductive values, defined as the relative contribution of individuals in a given class to the future of the population. These reproductive values are generally constant weights calculated from a constant projection matrix. Here, I show, for large populations and clonal reproduction, that reproductive values can be defined as time-dependent weights satisfying dynamical demographic equations that only depend on the average between-class transition rates over all genotypes. Using these time-dependent demographic reproductive values yields a simple Price equation where the non-selective effects of between-class transitions are removed from the dynamics of the trait. This generalises previous theory to a large class of ecological scenarios, taking into account density-dependence, ecological feedbacks and arbitrary distributions of the trait. I discuss the role of reproductive values for prospective and retrospective analyses of the dynamics of phenotypic traits.

**Note on this version:** Compared to the previous version of the manuscript, some changes have been made to improve the readability and structure of the text, and to clarify the connections with the existing literature. One reviewer pointed out inconsistencies in some of the numerical simulations, and in the mutation term in appendix A2, which have now been fixed. When checking the calculations, I have also identified a missing term in the equation for the dynamics of individual reproductive values. The new equation makes much more sense. Additional results are also presented, in particular for discrete-time models and populations with a continuous age structure, for which Fisher’s original definition of reproductive value can be recovered.

---

Evolution is fuelled by the genetic variance of populations. However, natural populations also display non-genetic sources of heterogeneity, when individuals of a given genotype belong to distinct classes representing different demographic, physiological or ecological states, with different demographic or ecological impacts on the population dynamics. This occurs for instance in an age-structured populations (e.g. when older individuals have a lower fecundity than younger individuals), in species with distinct developmental stages (e.g. when a species’ life cycle consists of a dispersing and a sessile morph), or in size-structured populations. The spatial location of an individual, or the quality of its habitat, may also be used to partition the population into distinct classes. In demography and ecology, this has led to a vast theoretical literature aiming at describing the population dynamics of such structured populations (Metz & Diekmann, 1986; Caswell, 2001).

In most theoretical analyses, intrinsic differences between classes of individuals are taken into account by weighting individuals by their reproductive values (Fisher, 1930; Price & Smith, 1972; Taylor, 1990; Rousset, 1999; Leturque & Rousset, 2002; Rousset, 2004; Rousset & Ronce, 2004; Engen et al., 2009; Engen et al., 2014). These reproductive values are defined as the long-term contribution of individuals in a given class to the future of the population, relative to the contribution of other individuals in the population. Reproductive values are typically calculated as a left eigenvector associated to the dominant eigenvalue of a constant projection matrix (Tuljapurkar, 1989; Taylor, 1990; Caswell, 2001; Rousset, 2004). Hence, the reproductive values are associated to the longterm growth rate of an exponentially growing population. Reproductive values play a key role in evolutionary game theory and inclusive fitness theory, where one seeks to compute the invasion fitness of a rare mutant arising in a monomorphic resident population that has reached its ecological attractor (Metz et al., 1992; Rousset, 2004; Metz, 2008; Gardner et al., 2011; Lehmann & Rousset, 2014). Under weak selection, the resulting selection gradient takes the form of a weighted sum of selective effects, where the weights are the class frequencies and the reproductive values calculated in the resident population (Taylor & Frank, 1996; Frank, 1998; Rousset, 1999; Rousset, 2004; Lehmann & Rousset, 2014; Gardner, 2015).

Reproductive values have also been used in combination with the Price equation (Price, 1970) in attempts to isolate the effect of natural selection from the effects of transitions between demographic classes (Crow, 1979; Engen et al., 2014; Grafen, 2015b). The motivation for doing so is the realisation that, in class-structured populations, the mean trait can change even in a neutral model where the vital rates do not depend on the trait, due to the dynamics of class structure itself. Following Grafen (2015b), I will refer to this latter effect as “passive changes”, to distinguish it from the effect of selection. In models with constant projection matrices, passive changes in mean trait are typically transient and disappear when a stable class structure is reached (reviewed in Tuljapurkar, 1989; Caswell, 2001). As first suggested by Fisher (1930), it is possible to get rid of this transient effect *from the start* if one uses reproductive values as weights when calculating the average phenotypic trait (Engen et al., 2014; Gardner, 2015). However, it is not clear how this property extends to models with density dependence or environmental feedbacks.

In this manuscript, I derive a class-structured Price equation coupled with a general ecological model in both continuous and discrete time. This extends previous works by Day & Gandon (2006) and Gandon & Day (2007), and gives an ecological underpinning to some results of Grafen (2015b). I then show, using only minimal ecological assumptions, that the purely demographic effect of class dynamics can be removed from the dynamics of the mean trait if one weights the mean trait in each class at time *t* by the reproductive value of that class at time *t*. This result is valid for large population sizes and clonal reproduction, but holds generally for any out-of-equilibrium ecological model, allowing for density-dependence, environmental feedbacks and environmental stochasticity. The requirement is that reproductive values are not calculated asymptotically in a population at equilibrium, but from a dynamical equation depending on *average* transition rates between classes, where the average is taken over all the genotypes. Related dynamical equations have been derived before in monomorphic populations (Tuljapurkar, 1989; Rousset, 2004; Rousset & Ronce, 2004; Barton & Etheridge, 2011), but to my knowledge their implications for the Price equation under general ecological scenarios have not been discussed. I discuss the usefulness of reproductive-value weighting for more practical studies, distinguishing between backward studies where one is interested in detecting selection in a known temporal series, and forward studies where one is interested in making predictions about the future change in a trait of interest. In particular, I show how these results extend previous results on the selection gradient calculated from traditional invasion analyses (Taylor, 1990; Metz et al., 1992; Taylor & Frank, 1996; Rousset, 1999; Rousset, 2004).

The structure of the paper should provide two levels of reading. Readers unfamiliar with the mathematical details behind the concept of reproductive value are encouraged to read up to section 3, where the potential usefulness of time-dependent reproductive values for analysing time series is discussed using numerical simulations. Sections 4 and 5 go deeper into the technical details and connections with previous studies and can be skipped at first read by non-theoreticians. The discussion should be readable by both categories of readers.

## 1 Ecological dynamics

The key points of the argument are easier to grasp using a population with a discrete structure and continuous-time dynamics. These assumptions will therefore be used in the primary derivation of the results, but extensions to discrete-time dynamics and continuous population structure will be discussed at a later stage. Table 1 provides a summary of the mathematical symbols used in this article.

I consider an infinitely large population, such that demographic stochasticity can be ignored. The population consists of *M* clonally reproducing types. A type may represent an allele or a phenotype, depending on the level of interest. The population is further structured into *K* classes. Throughout the article, I use the subscript *i* to refer to types and superscripts *j* and *k* to refer to classes. Hence, I denote the density of individuals in class *k* as *n*^*k*^ and the density of type *i* individuals in class *k* as 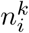 (the term density must be understood in the usual ecological sense, as number of individuals per surface area). These densities are collected in the vectors 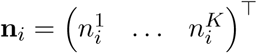 and 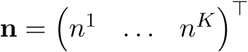.

Apart from clonal reproduction and large population densities, I will make only minimal ecological assumptions. The results are only expressed in terms of the transition rates *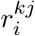* of *i* individuals from class *j* to class *k*. These transitions can be due to reproduction, mortality, maturation, or dispersal depending on the biological context. For instance, the production of class-*j* offspring by type-*i* parents in class *k* will contribute positively to the rate *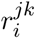*, while the death of type-*i* individuals in class *k* will contribute negatively to *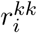*. Similarly, the movement of type-*i* individuals from class *k* to class *j* (due to maturation, dispersal, infection…) will add to *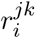* and subtract from *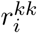* In general, the rates 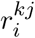 will depend on the vital rates of the focal type (fecundity, mortality, migration, infection…),but also on the vital rates of the other types. Most importantly, the rates *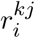* also depend on the environment **E**(*t*). The environment is defined from an individual-centred perspective (Metz et al., 1992; Mylius & Diekmann, 1995; Lion, in press) and collects all the relevant information necessary to compute the reproduction and survival of individuals. Basically, the vector **E**(*t*) collects the densities of the various types in the population, through the vectors **n**_*i*_, but also any ecological effects that are external to the focal population, which are collected in a vector **e**. These external effects may represent predation, parasitism, interspecific competition, or changes in abiotic factors. For spatially structured populations, other variables summarising the spatial distribution of types and individuals can also be added to the environment.

In continuous time, the dynamics of the total densities in each class may be written in matrix form as

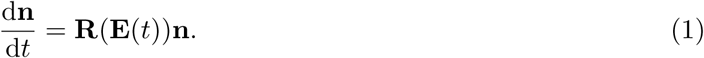

The matrix **R** has element 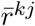 on the *k*th line and *j*th column, where 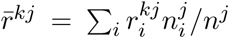 is the average transition rate from class *j* to class *k*. Coupled with a dynamical equation for the vector of external densities **e**, equation (1) forms the basis for ecological studies of class-structured populations (e.g. Caswell, 2001). For simplicity, I will often omit the dependency of the transition rates on the environment **E**(*t*) in the following, but it is important to keep in mind the generality of this formulation.

**Table 1:**
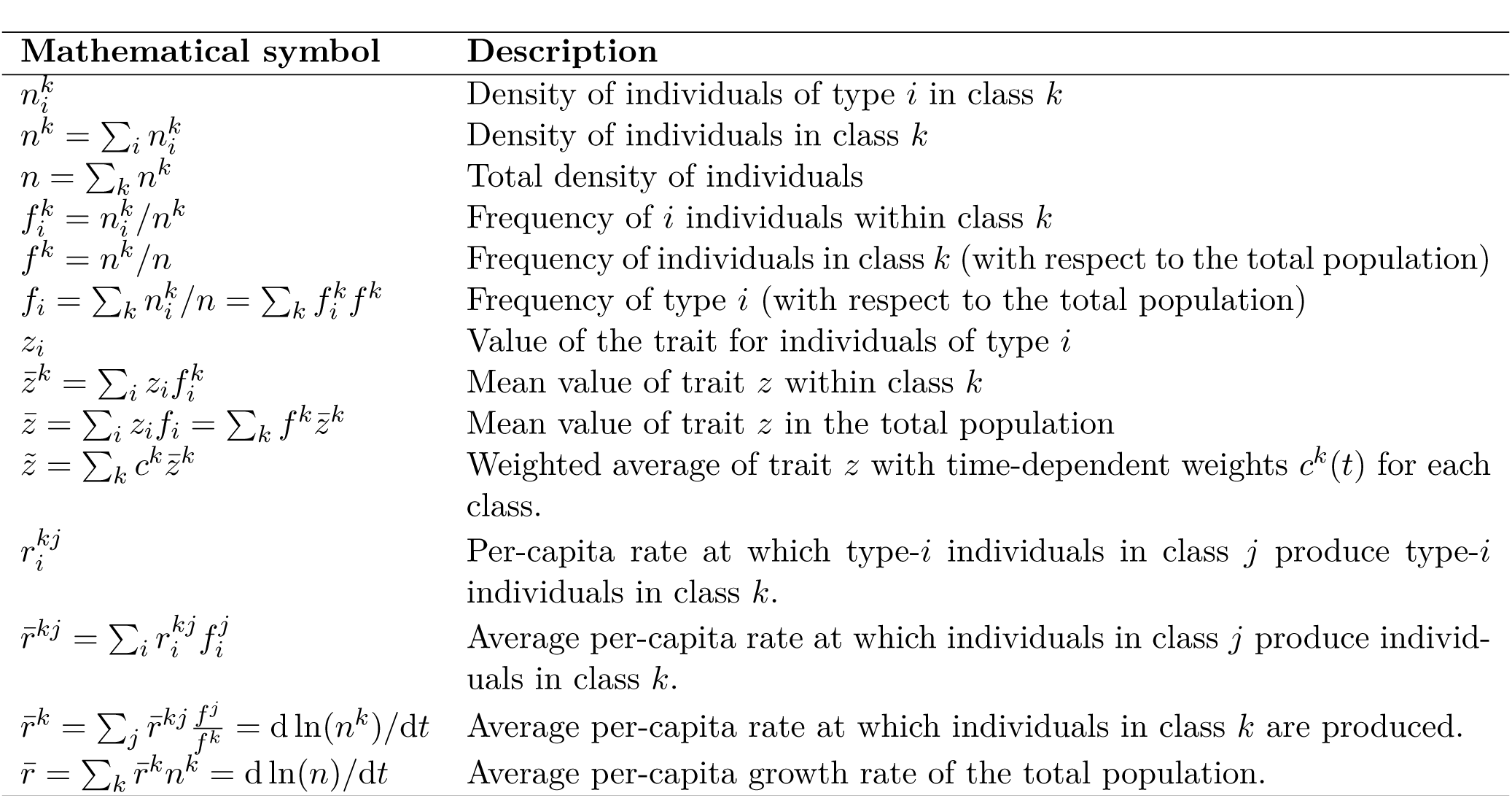
Definition of mathematical symbols used in the text

## 2 Dynamics of a phenotypic trait

Consider a trait *z*, with value *z*_*i*_ for type *i*. For simplicity, the trait is assumed to be measurable in each class and non-plastic (the trait value is fixed and does not depend on class, but see the discussion for how to deal with plastic traits). To study evolutionary change, I will focus on the change in the average of the focal trait, 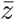. This average can be calculated in two equivalent ways, either direcly as 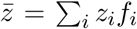, where *f*_*i*_ represents the frequency of type *i* in the population, or as a weighted sum of class means, 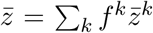, where 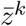 is the mean trait in class *k*, and *f* ^*k*^ is the frequency of class *k*.The frequencies of each class can be calculated as *f* ^*k*^ = *n*^*k*^*/n*, where *n* = *∑*_*k*_ *n*^*k*^ is the total density of individuals. The within-class average 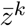 can be calculated as 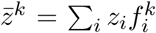, where 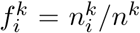 is the frequency of type *i* within class *k*. Throughout the manuscript, an overbar with a superscript will represent an average using the within-class frequencies *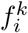* (as in 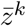), and an overbar without a superscript represents an average using the population frequencies *f*_*i*_ (as in 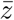).

### 2.1 The class-structured Price equation

In Appendix A, I show that the dynamics of 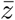 are given by the following differential equation,

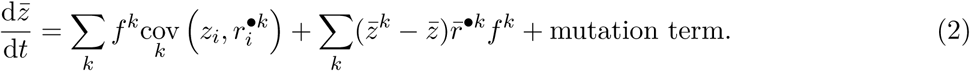

Equation (2) is the class-structured version of Price equation and shows that the change in mean trait can be partitioned into three components. The first term is the weighted average of the within-class covariances between the trait and the rate at which each individual of type *i* in class *k* produces individuals in any class, 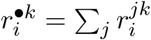, which is measure of fitness of type-*i* individuals in class *k*. By definition, 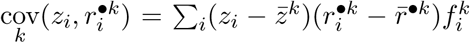 The second term is the between-class covariance between the mean trait in a class and the average rate at which an individual in class *k* produces individuals in any class, 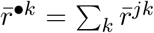. This term depends on the phenotypic differentiation between a given class and the total population, 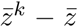. Hence, equation (2) partitions the change in mean trait into a within-class and a between-class component. Finally, the third component of equation (2) represents the effect of mutation, recombination, or possibly external immigration events. In the following, I will neglect the mutation term and focus on the effects of natural selection and demographic changes on the dynamics of the mean trait, but a more complete description of the mutation term can be found in Appendix A.

Now let us assume that the per-capita growth rates 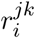 are independent of the type (i.e. 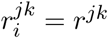 for all *i*). Intuitively, we should not observe any selection in such a population. Indeed, equation (2) tells us that the covariances in the first term are zero. However, one might still observe directional change in the mean trait due to the second term, which depends on the average rates 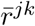 and not on the correlation between the trait and the type-specific transition rates. Following Grafen (2015b), I will refer to this effect as the “passive changes in mean trait”.

Passive changes in mean trait obviously disappear if the class means exactly coincide with the population average. The mechanisms causing the build-up of between-class differentiation can be elucidated by writing the equation giving the dynamics of the mean trait in class *k*, 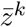 (Appendix A). Dropping the mutation term for simplicity, this gives:

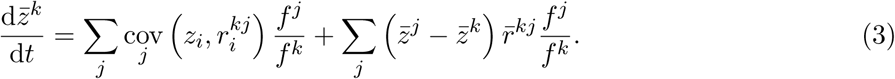

This shows that there are two components driving the dynamics of between-class differentiation. Even when the per-capita growth rates 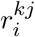 are independent of the trait, so that the covariance terms are zero, the mean trait within class *k* can still change due to between-class demographic transitions between class *k* and the other classes. This can lead to changes in the phenotypic differentation across classes, measured by the deviation of the class averages 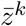 from the population average 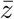 Hence, the second term of equation (2) conflates the consequences of natural selection (through the covariance term in equation (3)) and of other ecological or genetical mechanisms causing phenotypic differentiation between classes.

### 2.2 The class-structured Price equation for a weighted average

Equation (2) is derived by giving each individual a weight of unity. In contrast, a common approach in the literature has been to assign a class-specific weight to each individual in order to extract the signal of natural selection from the change in mean trait (Fisher, 1930; Crow, 1979; Taylor, 1990; Taylor & Frank, 1996; Leturque & Rousset, 2002; Rousset, 2004; Rousset & Ronce, 2004; Engen et al., 2014; Grafen, 2015b). Here, I follow this approach and consider the dynamics of a weighted average of the focal trait. In contrast with the standard practice, however, I consider that the individual weights are not constant through time. I therefore give each individual in class *k* at time *t* a weight of *v*^*k*^(*t*). A weighted average for the focal trait can then be calculated at time *t* as

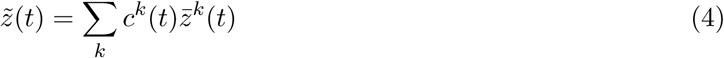

where the weight *c*^*k*^(*t*) = *v*^*k*^(*t*)*f*^*k*^(*t*) is assigned to class *k* at *t* and scaled such that ∑_*k*_ *c*^*k*^(*t*) = 1. When all the *v*^*k*^’s are set to the constant value 1, we recover the results of the previous paragraph. With these assumptions, the change in the weighted mean trait takes the following simple form:

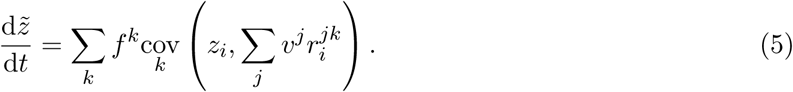

if the weights *c*^*k*^(*t*) satisfy a particular dynamical equation (Appendix A). Hence, for a well-chosen set of weights, we can write the change in mean trait as the average across all classes of the covariance between the trait and the (weighted) mean contribution of individuals in that class. The change in a neutral trait with no effect on the vital rates will therefore be exactly zero. Comparing the covariance term in equation (5) to the covariance term in equation (2), we note that the only difference is that the sum *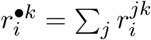* is replaced with the weighted sum 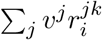

For equation (5) to hold, the *c*^*k*^’s must satisfy the following system of differential equationsFor equation (5) to hold, the *c*^*k*^’s must satisfy the following system of differential equations

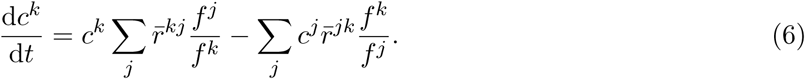

This equation takes the form of a master equation describing the time evolution of a vector of proabilities. An interpretation of this equation will be given in the next section.

#### Discrete-time dynamics

For comparison with other works, it is useful to consider the discrete-time version of this result. In Supporting Information S.1, I show that the change in weighted mean trait can be written in discrete time as

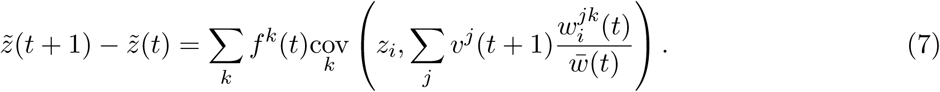

Compared to equation (5), the per-capita growth rates *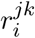* are replaced by the relative fitnesses 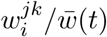 where 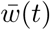 is the average fitness in the population and the class weights satisfy the following recursion:

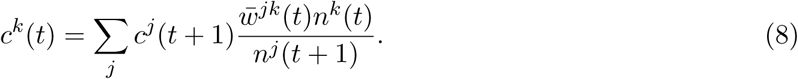

which is a discrete-time analog of equation (6).

Importantly, the elimination of passive changes holds if the ck’s satisfy equation (6) and (8), irrespective of initial or final conditions. As a result, the vector of weights is not unique, and additional considerations are required to choose the relevant final condition. I will come back to this point when presenting the numerical applications of this approach.

### 2.3 Biological interpretation of the weighted average

So far, the validity of the results does not hinge on a particular biological interpretation of the weights used to compute the average 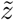 However, it turns out that the weights *v*^*j*^(*t*) that appear in equation(5) are the individual reproductive values in each class at time *t*, and that the weights *c*^*j*^(*t*) are the class reproductive values at time *t* (Taylor, 1990; Rousset, 2004).

Indeed, a biological interpretation of *c*^*k*^(*t*) can be given as the probability that a random gene sampled at some time in the future has its ancestor in class *k* at time *t* when we look backward in the past. In discrete time, this probability will satisfy the following recursion

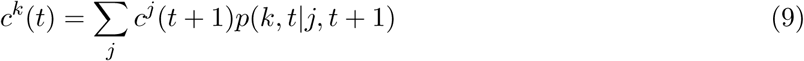

where *p*(*k, t| j, t* + 1) is the probability that the lineage was in class *k* at *t* given that it is in class *j* at *t* + 1 (see e.g. Rousset (2004), equation (9.21)). This probability is the fraction of class-*j* offspring produced by class-*k* individuals at time *t*, which is simply the number of class-*j* offspring produced by class-*k* parents, 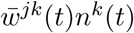 divided by the total number of class-*j* offspring, *n*^*j*^(*t* + 1). Hence, equation (9) is exactly equation (8).

A perhaps more intuitive, but equivalent, interpretation of *c*^*k*^(*t*) can be given as the (relative) number of descendants left by genes present in class *k* at time *t*, from *t* onwards (Tuljapurkar, 1989; Caswell, 2001; Rousset, 2004; Barton & Etheridge, 2011), which is exactly the definition o f reproductive value as a measure of relative long-term contribution used in population genetics and demography (going back to Fisher (1930) and Goodman (1968)). The connection with previous definitions o f reproductive values will be explored in more detail in section 4.

The previous analysis thus shows that the individual reproductive values can be used as time-dependent individual weights that guarantee the elimination of the passive changes in mean trait at any time. The discrete-time formulation (equation (8)) more clearly shows that the reproductive-value weighting needs to be applied to the offspring generation: an offspring in class *j* (at generation *t* + 1) is valued by its current contribution to the future of the population, *v*^*j*^(*t* + 1).

In continuous time, the distinction between parent and offspring generations is blurred, but equation (6) can still be interpreted in a similar way. Indeed, because the terms for *j* = *k* cancel out, the first term on the right-hand side is 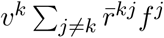 T his tells us that the probability for a gene lineage to be in class *k* at time *t* will increase due to the creation of new class-*k* individuals, with reproductive value *v*^*k*^, from individuals belonging to all the other classes. The second term is 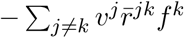 and shows that the probability for a gene lineage to be in class *k* at time *t* will decrease due to the creation of new class-*j* individuals, with reproductive value *v*^*j*^, from class-*k* individuals. It can be shown that the master equation (6) is the continuous-time limit of the discrete-time backward recursion (8).

## 3 Reproductive values for retrospective data analyses

In this section, I present numerical simulations to show how the dynamical reproductive values can be computed from time series and used to remove the passive changes from the dynamics of the mean trait. As a proof-of-concept, I consider a discrete-time three-class model, with class densities *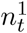*, 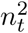 and 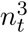. The transition matrix for type *i* at time *t* is given by

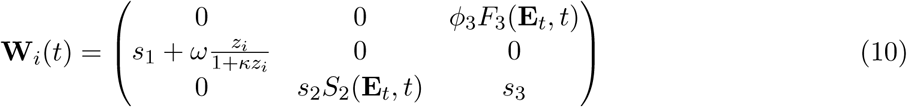

The elements of **W**_*i*_ are the 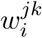 of equation (7). The model is a variation on the classical Larva-Pupae-Adult (LPA) model for the dynamics of *Tribolium* populations (Dennis et al., 1995). The reproduction and survival of stages 2 (pupae) and 3 (adults) depend on the environmental dynamics through the fecundity function *F*_3_(**E**_*t*_*, t*), and the survival function *S*_2_(**E**_*t*_*, t*), for which I make the following assumptions:

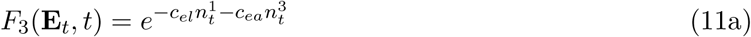

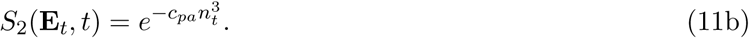

Following traditional notation, *c*_*el*_ (resp. *c*_*ea*_ and *c*_*pa*_) reflects the intensity of cannibalism of eggs by larvae (resp. eggs by adults and pupae by adults). Individuals are characterised by a trait *z*, which is a property of the type and confers a non-linear survival advantage to the first stage. The parameter *ω* measures the strength of selection.

### General method

Starting from some initial conditions, the model can be run forward in time from time 0 to time *T* to provide a sequence of data. The details of the model are irrelevant, but what matters in the end is that we get a time series for the mean traits in each class, 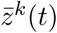 for the class densities *n*^*k*^(*t*), and for the average fitnesses 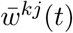, which determine between-class transitions. These quantities can in principle be measured in the field without any knowledge of the genetic variation in the population. Knowing this, recursion (8) can be iterated backward in time, starting from a given final condition **c**(*T*), yielding the weights **c**(*t*) that need to be applied to the mean traits 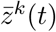 at each time step. I will first present two illustrating examples, before discussing the choice of the final condition.

### Example 1

Figure 1a shows that, even for a neutral trait (*ω* = 0), the model exhibits sustained fluctuations in the mean trait. A naive observer may interpret these fluctuations as caused by selection, but in fact this is only due to demographic transitions between classes. The absence of selection is revealed by plotting the dynamics of the weighted mean trait, using the class reproductive values computed from equation (8) as weights. Doing so gives a flat line (in red), which reveals that the fluctuations are not caused by selection. For comparison, the lower panels of figure 1b also show the dynamics of the mean trait weighted with constant reproductive values calculated from the time-averaged projection matrix (gray line). This weighting does not completely remove the passive changes in mean trait. In the model with selection (figure 1b), the mean trait appears to fluctuate around an increasing trend. Applying our reproductive-value weighting irons out the fluctuations due to the passive changes in mean trait and yields a smooth trajectory that reveals the part of the change in mean trait that is actually due to selection. Again, using constant reproductive values calculated from the time-averaged projection matrix does not eliminate the passive changes in mean trait (gray line). Using the time-dependent neutral reproductive values from figure 1a as weights (blue dashed line) also imperfectly removes the passive changes, even though selection is assumed to be weak in the model.

**Figure 1:**
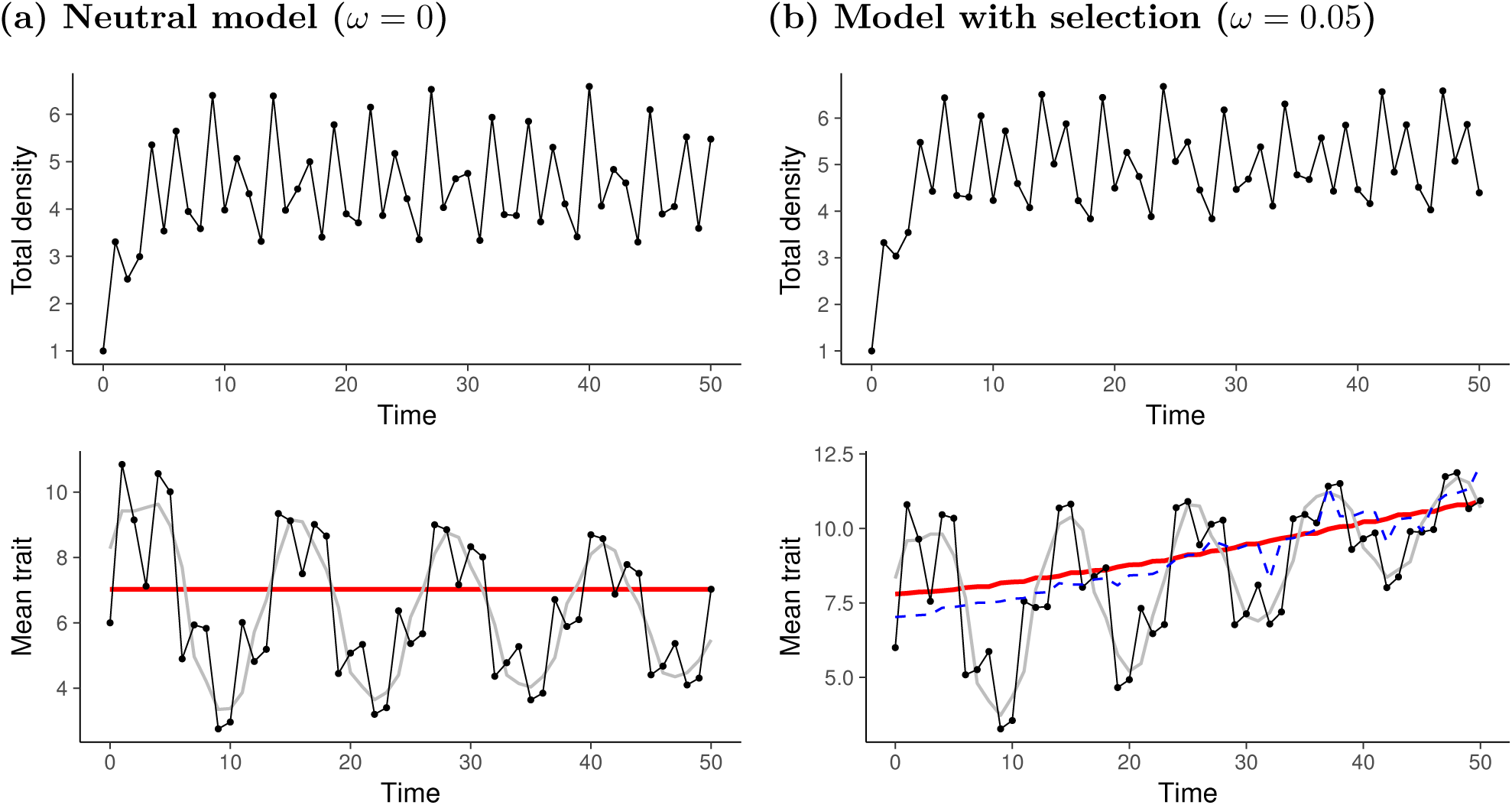
The dynamics of total population density and mean trait are shown for Model 1 at neutrality (left panel) and in the presence of selection (right panel). The dynamics of the model are given by equation **n**_*i*_(*t* + 1) = **W**_*i*_(*t*)**n**_*i*_(*t*), where the transition matrix is defined in equation (10) and (11). The upper panel gives the dynamics of the total population size, *n*(*t*). The lower panel gives the dynamics of the arithmetic mean of the trait (dots), of the reproductive-value weighted trait (red line), and of the weighted mean trait using constant reproductive values computed from the average matrix over time (gray line). In figure (b), the blue dashed line shows the dynamics of the weighted mean trait using the neutral reproductive values.. The initial densities for each class are *n*_1_(0) = 0.3, *n*_2_(0) = 0.3, *n*_3_(0) = 0.4. The initial distribution of the types is Poisson with means 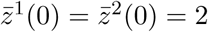 and 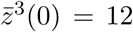, so that 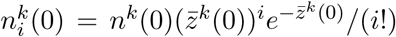 and *zi* = *i* for 0 *≤ i ≤* 49. Parameters: *φ*_3_ = 10, *s*_1_ = 0.6, *s*_2_ = 1, *s*_3_ = 0.05, *c*_*ea*_ = 0.5, *c*_*pa*_ = 1, *c*_*el*_ = 0.4.

### Example 2

I now assume that fecundity in stage 3 is further affected by environmental stochasticity, through a stochastic multiplicative factor *ρ*_*t*_. Figure 2 shows that, even with environmental stochasticity, reproductive-value weighting can eliminate the transient passive changes in mean trait. Note that, with environmental stochasticity the transient fluctuations decay rapidly and, eventually, the weighted and unweighted averages of the trait coincide.

**Figure 2:**
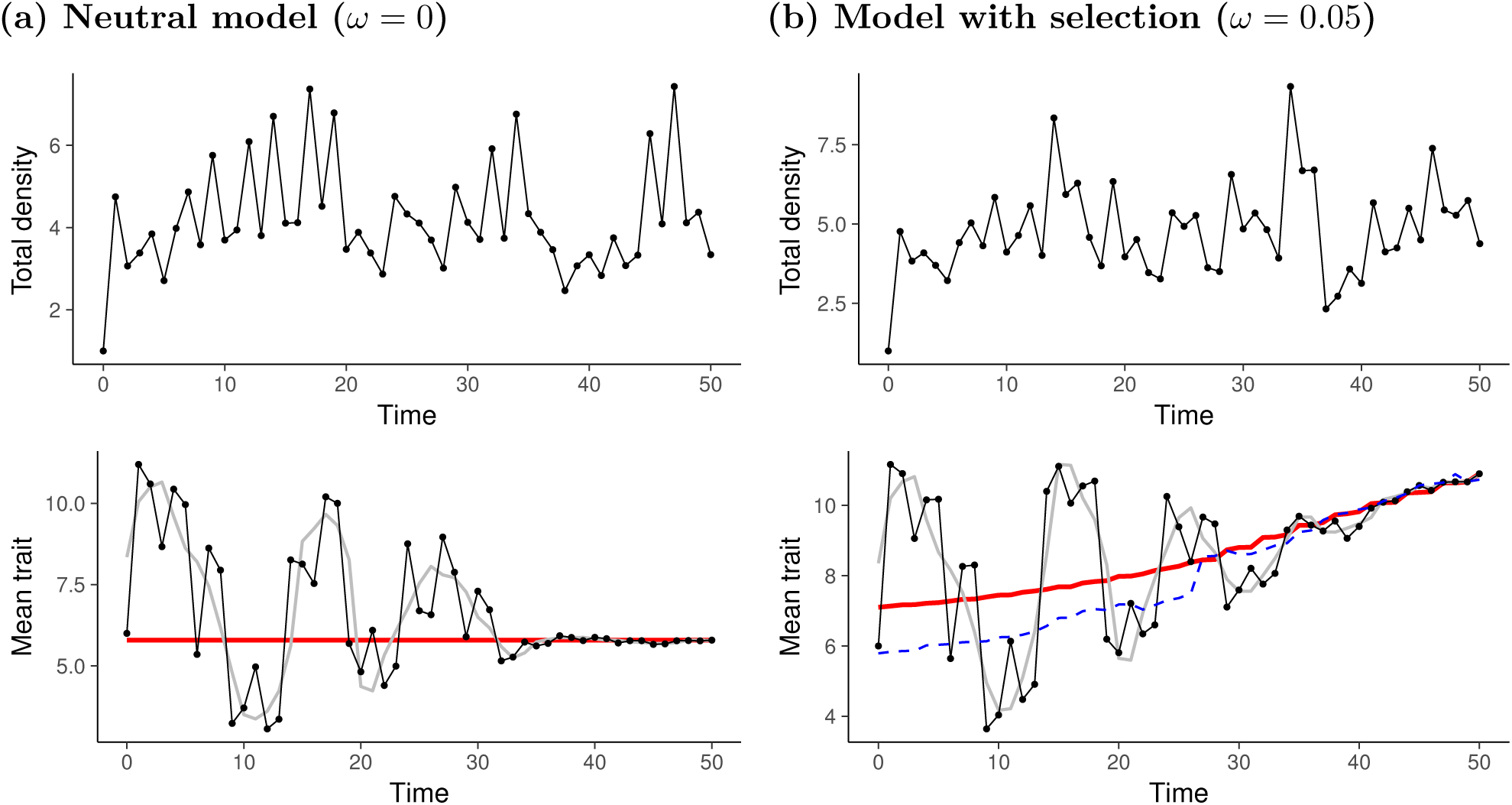
The dynamics of total population density and mean trait are shown for Model 2 at neutrality (left panel) and in the presence of selection (right panel). Compared with Model 1 and figure 1b, the only change is that the fecundity *F*_3_(**E**_*t*_*, t*) in the matrix **W**_*i*_(*t*) is multiplied by a stochastic factor *ρ*_*t*_, modelled as a uniformly distributed random variable between 0.5 and 1.5. To allow comparison, the same sequence of random numbers is used in figure 2a and 2b.

### Choice of the final condition

As noted above, the choice of the final condition is irrelevant when deriving equations (5) and (7). In fact, for a neutral trait, the dynamics of the weighted mean trait should be a flat line, irrespective of the final condition. With selection, however, different final conditions will yield different trajectories for the weighted trait. In the two examples above, I used the final condition **c**(*T*) = **f** (*T*) to compute the class reproductive values and weighted mean trait at each time. The choice of this particular final condition is equivalent to setting the relative contribution of each individual to the present generation to 1 (Barton & Etheridge, 2011), but also guarantees that the trajectory of the weighted mean trait converges to the value measured at the end of the time series. In other words, from the final state of the population under study, we trace backward in time the trajectory corresponding to the change in mean trait in an ideal population where the passive changes have been removed.

A further motivation for choosing this final conditions comes from the consideration of the limiting regime where selection is weak. The influence of the passive changes in mean trait should decay rapidly under weak selection. As a result, if we have enough data points, we can expect the weighted dynamics to converge to those of the unweighted mean trait, as in figure 2b.

## 4 The interplay of demography and selection

The previous results show that the effect of selection in class-structured populations is best captured by weighting each class with time-dependent reproductive values. Using this weighting yields a compact expression for the dynamics of mean phenotypic traits, (5), which can also be written in matrix form as follows

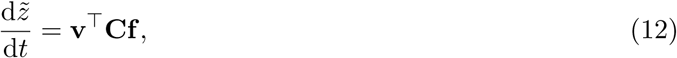

where **C** is the matrix of covariances with components 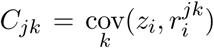. In the remainder of the paper, the notation *┬* represents the transpose operation, i.e. **v**^*T*^ is a row vector.

### 4.1 Dynamical equations for the demographic process

The vectors of reproductive values and class frequencies follow coupled dynamical equations that generalise the classical interpretation in terms of eigenvectors. For class reproductive values, equation may be written compactly in matrix form as

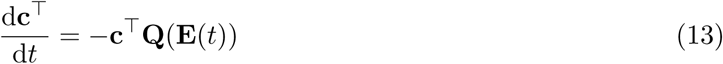

where **E**(*t*) is the vector of environmental variables and **Q**(**E**(*t*)) is the matrix with elements *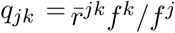* for *j* ≠ *k* and *q*_*kk*_*=-∑* _*j* ≠ *k*_ *q*_*kj*_(Appendix B).

Similarly, the vectors **v**^*┬*^ and **f** in equation (12) satisfy the following equations (Appendix B)

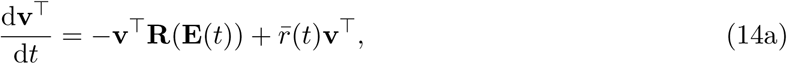

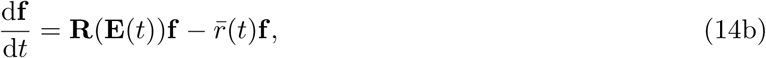

where 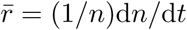 is the average growth rate of the total population. Note the similarity between these two equations. The individual reproductive values and class frequencies are both calculated using the matrix **R**(**E**(*t*)), which collects the average transitions rates 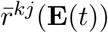, and the per-capita growth rate of the total population size, 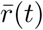. However, while equations (14b) and (14a) use the same input, they have different interpretations: using equation (14b), the future class frequencies can be calculated from an initial condition, whereas equation (14a) works backwards in time to compute past reproductive values from a final condition.

Equations (14b) and (14a) can be combined to recover an important property of the individual reproductive values already noted by Fisher (1930) for linear models. If we evaluate the population size not by a head count, but by weighting each individual by its reproductive value, the weighted population size, 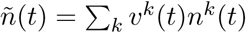 satisfies the following equation

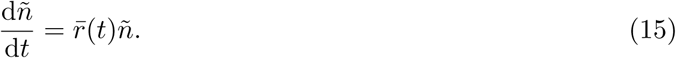

Hence, the reproductive value-weighted population size always grows as the unweighted population size, even for out-of-equilibrium, non-linear ecological dynamics.

Note that it is also possible to derive similar equations for populations with continuous structure. I give an illustration in Box 1 using a model with continuous age structure which allows to recover Fisher’s original definition of reproductive value.

#### Discrete-time dynamics

Equations (13) and (14a) have discrete-time counterparts. In Supporting Information S.1, I show that the class reproductive values satisfy the following recursion:

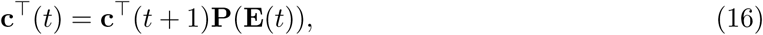

where **P** is the matrix with elements 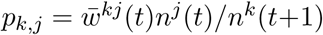. Similarly, the individual reproductive values and class frequencies satisfy the recursions

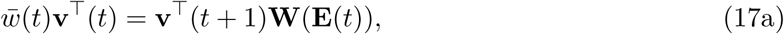

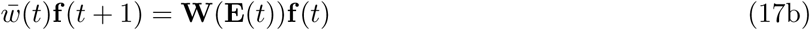

where **W**(**E**(*t*)) is the matrix with elements 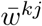. Again, the two equations use the same input, but represent different processes. As already noted by Tuljapurkar (1989), the matrix **W** acts to propagate **f**^*T*^ forward in time and **v**^*T*^ backward in time. In Tuljapurkar’s words, the vector of class frequencies at each time is an “accumulation of the past”, while the vector of reproductive values at each time is a “summation of the future”.

### 4.2 Selection and demography

Equation (12) provides a simple partition of the change in the weighted mean phenotype into selective and demographic components. While the matrix of covariances **C** captures the relative performances of the different types for each between-class transition, the vectors **v**^T^ and **f** solely depend on the average rates at the population level and are therefore purely demographic properties of the system. Indeed, a naive ecologist oblivious to the underlying genetic diversity of her study population would still be able to calculate the reproductive values and class frequencies from the aggregate response of the population. This is potentially valuable for data analysis, as we have seen in the previous section.

This demographic definition of reproductive values has its roots in Fisher (1930)’s original exposition of the concept and provides a conceptually clear connection to the usage of reproductive values in other fields, such as human demography, where details about the genetic composition of the population are typically averaged out. Importantly, the dynamical definition of reproductive values used here holds for a broad class of models, irrespective of the genetic composition of the population, of the trait distribution, and of the underlying population and environmental dynamics. In particular, the reproductive values are not calculated in a neutral or monomorphic population, nor under any limiting assumption of mutant rarity, as typically assumed in evolutionary game theory. The next paragraph clarifies the connection with these previous usages of reproductive values.

### 4.3 Connection with classical asymptotic results

Although most works directly compute reproductive values as a left eigenvector, some authors have proposed to compute reproductive values from dynamical equations. To my knowledge, a dynamical equation was first proposed by Crow (1979) for allele-specific reproductive values and by Tuljapurkar (1989) at the population level (see also Barton & Etheridge (2011)). Dynamical equations for the class reproductive values in monomorphic populations have also been used in inclusive fitness theory (e.g. equation 9.21 in Rousset (2004) or equation (6) in Lehmann (2014)). However, while Tuljapurkar (1989) explicitly defined reproductive values as a function of time, usage in evolutionary theory has typically reserved the word “reproductive value” for the asymptotic behaviour of the dynamical equations, yielding a time-independent definition (Charlesworth, 1994; Rousset, 2004; Barton & Etheridge, 2011; Lehmann, 2014). This asymptotic definition of reproductive values hinges on additional demographic or genetic assumptions, such as exponential growth or weak selection, although it has been noted that, in principle, reproductive values could be defined as time-dependent weights, as I do here (see Lehmann & Rousset (2014), note 3). In this section, I discuss how previous definitions of reproductive values can be recovered from equations (13) and (14a) under additional assumptions.

#### 4.3.1 Exponential growth

The easiest way to recover the standard asymptotic definition of reproductive values is to assume that the matrix **R** is approximately constant^1^. Then, it is well known that, in the long run, the population growth rate 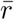 is constant and equal to the dominant eigenvalue of **R**. The population then grows exponentially with a stable class structure given by the right eigenvector of matrix **R** associated to 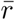 The corresponding left eigenvector collects the individual reproductive values (Goodman, 1968; Tuljapurkar, 1989; Caswell, 2001). Reproductive value can then be defined as this left eigenvector, which gives the long-term contribution of individuals in a given class to the future of the population, relative to the contribution of other individuals in the population.

Equations (14) allow us to recover this result by assuming that the reproductive values and class frequencies stabilise in the long run. Setting the time derivatives to zero then yields

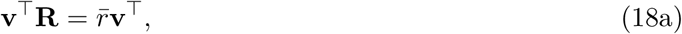

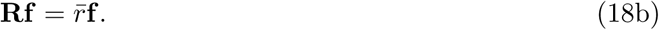

The analog discrete-time result follows from setting **v**(*t* + 1) = **v**(*t*) and **f** (*t* + 1) = **f** (*t*) in equation (17), which gives

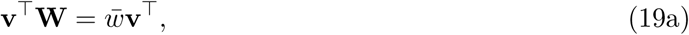

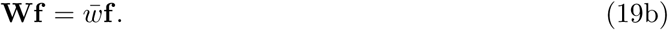

The latter equations are respectively the leftand righ-eigenvector results of Taylor (1990) (his equations (7) and (5)), but they do not rely on the assumption that the population is monomorphic. Rather, they explicitly take into account polymorphic populations with arbitrary trait distribution. The key to this generalisation is to use the matrix of average transition rates.

#### 4.3.2 Density-dependent populations at equilibrium

Another frequent assumption in the literature is that the population is at a stable demographic equilibrium. Then, the dynamics of reproductive values also depend on constant projection matrices 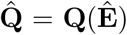 and 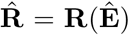, where the environmental vector 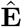 is calculated at equilibrium. Because 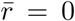 at equilibrium, it follows from equations (13) that the vector **c** is a left eigenvector of the matrix 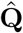 associated to eigenvalue 0, where 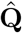 has elements 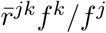. Similarly, equations (14a)-(14b) show that the individual reproductive values are proportional to a left eigenvector of 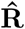 associated to eigenvalue 0, and the class frequencies to a right eigenvector.

A similar result holds in discrete-time. From equation (16), we see that at equilibrium when **c**(*t* + 1) = **c**(*t*) and 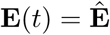, the vector **c** is a left eigenvector of the matrix 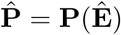 associated to eigenvalue 1. For the individual reproductive values, we have 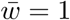 at equilibrium and therefore **v** is a left eigenvector of the matrix 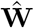 associated to eigenvalue 1. These two results extend a widely used result in evolutionary game theory and inclusive fitness theory (Taylor, 1990; Rousset, 1999; Rousset, 2004). Once again, in contrast to these previous studies, the reproductive values and class frequencies are defined in polymorphic populations with arbitrary trait distribution, instead of being calculated in a monomorphic population. However, it is straightfoward to recover the standard monomorphic case from the polymorphic population under the assumption that all types are identical.

## 5 Reproductive values for predictive theoretical analyses

### 5.1 Separation of time scales

By construction, reproductive values quantify class contributions to the future demography of the population. Equations (13)-(14a) and (16)-(17a) show that they can be calculated from backward dynamical equations. As a result, equation (12) appears to have little predictive power as the change in the mean trait at a given time depends on the whole future we are precisely trying to predict. However, this problem can be solved if we are only interested in long-term evolution and assume a separation of time scales between evolutionary and ecological time scales, as is typical when computing invasion fitness (Metz et al., 1992; Geritz et al., 1998; Lehmann & Rousset, 2014; Van Cleve, 2015). If evolution is slow compared to the demography of the population, we only need to evaluate equation (12) on the population’s ecological attractor, which can be a point equilibrium, a limit cycle, or more complicated objects. On the ecological attractor, the future is predictable, and the reproductive values give information about the long-term contribution of each class, as required to analyse long-term evolution.

Equation (12) can then be used as a starting point to derive approximations for the change in mean trait following the introduction of a mutation with small phenotypic effect. I will illustrate this idea in the remainder of this section.

### 5.2 Monomorphic resident population at equilibrium

I will first recall a classical result of evolutionary game theory obtained under the assumption of a vanishingly small trait variance in the population. Consider two types *w* and *m* with traits *z*_*w*_ and *z*_*m*_ = *z*_*w*_ + *ε*. When *ε* = 0, we assume that the population settles on a demographic equilibrium. For small values of *ε*, the following approximation for the change in weighted mean trait can be derived (Appendix S.2; Taylor (1990), Taylor & Frank (1996), and Rousset (2004))

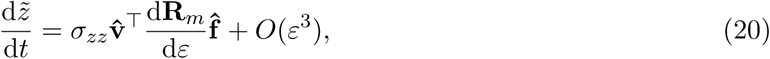

where the vectors 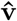 and 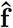 equation (20) are the equilibrium values of **v** and **f** computed in the monomorphic resident population, and the matrix d**R**_*m*_*/*d*ε* is the perturbation of the matrix of the mutant per-capita growth rates 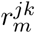 (see Appendix S.2 for more details). Note that 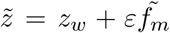, where 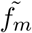 is the average frequency of the mutant type, calculated using class reproductive values as weights (as defined in Appendix A.3), so that tracking the average phenotype is equivalent to tracking the frequency of the mutant allele.

##### Box 1: Continuous age structure and Fisher’s original concept of reproductive value

Fisher (1930) originally defined the concept of reproductive value in a model with continuous age structure, whereas the results in the main text are derived using a discrete class structure. However, the elimination of passive changes through reproductive-value weighting also extends to models with continuous structure. Defining the reproductive value of individuals with age *a* at time *t* as *v*(*a, t*), it is possible to derive the dynamics of a weighted average trait, 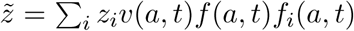, where *f*_*i*_(*a, t*) is the fraction of type-*i* individuals among individuals with age *a* at *t*, and *f* (*a, t*) is the fraction of individuals with age *a* at time *t*. In Appendix S.3, it is shown that the change in mean trait then takes the following form

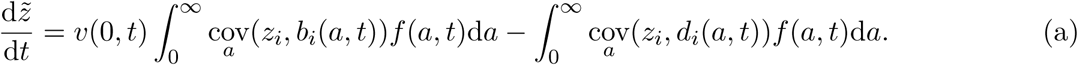

This is the continuous-age equivalent of equation (12). Here, *b*_*i*_(*a, t*) and *d*_*i*_(*a, t*) are the birth and death rates of type-*i* individuals with age *a* at time *t*. The covariances between the trait *z* and the vital rates are taken over all individuals with age *a* and time *t*. The contribution of the reproduction of individuals with age *a* to selection is weighted by their frequency *f* (*a, t*) and by the reproductive value *v*(0*, t*) of their newborn offspring. In contrast, the contribution of death events to selection is weighted by the reproductive value of the age group, *v*(*a, t*).

Equation (a) only holds if the reproductive values *v*(*a, t*) satisfy the following partial differential equation

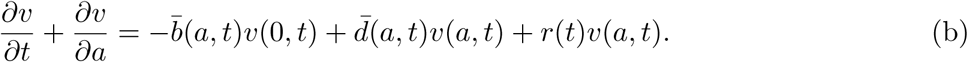

Again, this is the continuous analog of equation (14a). The reproductive values depend on the average birth and death rates 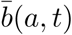 and 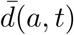 and on the growth rate of the total population 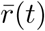. Bacaër & Abdurahman (2008) derived a similar equation in a monomorphic epidemiological model structured with infectious age.

To fully connect these results to Fisher (1930)’s original definition of reproductive values, one needs to assume time-independent birth and death rates that only depend on age. With these assumptions, the population will be characterised by a stable growth rate *r*, a stable age structure *f* (*a*) and a stable distribution of reproductive values *v*(*a*). From equation (b), we then have the following expression:

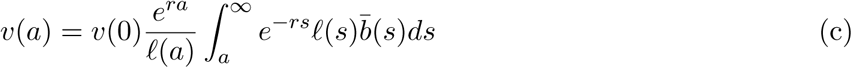

where *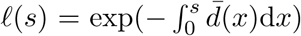* is the probability of surviving up to age *s*. Expression (c) is Fisher’s original expression of reproductive value (Fisher, 1930; Charlesworth, 1994), but it explicitly takes into account genetic polymorphism in the population by using the average birth and death rates.

Equations (12) and (20) have the same form, but the second is only valid as an approximation under weak selection. Expanding the matrix product in equation (20) then yields the classical expression for the selection gradient as a weighted sum of the effects of selection on class transitions (Taylor, 1990; Rousset, 1999; Rousset, 2004),

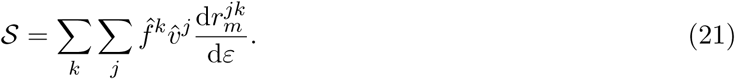

Selection gradients of this form are frequenly encountered in the literature, when quasi-monomorphic populations are considered. Quasi-monomorphism typically arises in two-allele models when the mutant allele is rare compared to the resident allele (as in Taylor, 1990), or in models with a continuous trait distribution, when the trait distribution is assumed to be tightly clustered around the mean (weak selection). Under these assumptions, the effect of a mutation on the demography of the population can be neglected compared to the effect of the mutation on the covariance matrix. The fundamental reason is that, because the resident population is monomorphic, **C** = **0** when *ε* = 0. As a result, the perturbations of the vectors **v** and **f** will contribute terms of higher order to the dynamics of the mean trait compared to the perturbation of the matrix **C**, and can therefore be neglected.

### 5.3 Polymorphic resident populations at equilibrium

Although standard models tend to focus on monomorphic resident populations, many natural populations will typically display a non-negligible amount of standing variation, with potentially multimodal trait distributions. Because equation (12) is still valid under these assumptions, it can be used as a starting point to derive approximations of the selection gradient. As in the monomorphic case, the idea is to calculate perturbations of equation (12) resulting from a slight change in the trait distribution. For instance, if we have a stable coalition of *M* types, we may consider that a fraction *p* of the individuals of type *M* mutates to trait value *z*_*M*_ + *ε*. As before, the limit *ε* = 0 corresponds to the resident population at equilibrium. If we can further assume that the effect of the mutation on the population demography is negligible compared to the perturbation of the covariance matrix **C**, we can approximate equation (12) as

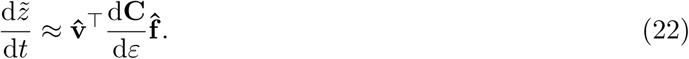

As in equation (20), the vectors 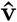 and 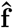 are computed at equilibrium for *ε* = 0. However, equation (22) is valid for arbitrary trait distributions in the resident population. The reproductive values and class frequencies must therefore be computed from the mean demographic matrix of the resident population, which is the natural extension of the “neutral” reproductive values typically considered when the resident population contains only one type.

Of course, additional work is needed to investigate the domain of validity of this approximation, which is far beyond the scope of this paper. In particular, because in the resident polymorphic population the covariance matrix **C** is not null, it may not always be possible to neglect the effect of the mutation on the demographic variables **v** and **f** (Appendix S.2). However, the present considerations shed light on the potential utility of equation (12) for deriving analytical expressions for long-term measures of selection in polymorphic class-structured populations, while keeping the central concept of reproductive value on board.

### 5.4 Periodic ecological attractor

Another potential extension of standard theory attainable from equation (12) is to consider nonequilibrium ecological attractors, such as limit cycles. Limit cycles can be thought of as a continuous-time description of periodic environments, as needed for instance for taking into account seasonality.

Consider a monomorphic population that has settled on a limit cycle with period *T*. Assuming as in the equilibrium case that selection is weak, the average change in the mean trait over one period is approximately proportional to (Appendix S.2)

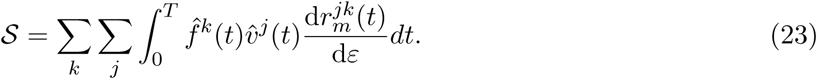

The reproductive values and class frequencies are time-dependent and computed using the matrix 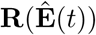, where 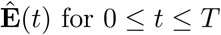 is the periodic environment generated by the resident population. The rates 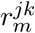 are also calculated on the resident environment, which is indicated by the dependency on *t*. The use of time-dependent reproductive values for periodic models has been suggested before for continuous-time exponentially growing populations (Bacaë r & Abdurahman, 2008) and discrete-time density-regulated populations (Brommer et al., 2000) but to my knowledge equation (23) has not been previously derived. Compared to earlier approaches that have dealt with complex demographies by incorporating the demographic states into the class-structure (Brommer et al., 2000; Rousset & Ronce, 2004; Lehmann et al., 2016), equation (23) provides a lower-dimensional invasion criterion in which classes are defined independently of the population dynamical model. For instance, if we study an ecological model with different attractors depending on parameter values, we do not need to change the class structure and the dimension of the projection matrix to analyse the different regions of parameter space. Although a full analysis of the connections between this result and previous characterisations of invasion fitness in periodic environments (Tuljapurkar, 1985; Ferrière & Gatto, 1995) is beyond the scope of this article, this preliminary attempt suggest that equation (12) could be used to provide potentially useful approximations for the change in mean trait also in non-stationary ecological systems.

## 6 Discussion

In class-structured populations, changes in gene frequencies or mean phenotypes can be brought about through three distinct routes. First, natural selection can act within each class through the covariance between the focal trait and the vital rates of each type within that class. Second, directional changes in the mean trait can occur due to the dynamics of between-class differentiation, as measured by the difference between the mean trait in a class and the mean trait in the total population. The dynamics of between-class differentiation is itself the resultant of natural selection and of “passive changes” due to transitions between classes. These passive changes can be observed even in the absence of natural selection, either transiently or on longer time scales, depending on genetic constraints and environmental feedback. Third, mutation or recombination may introduce some directional change in the mean trait, an effect that I have ignored in this article and should be kept in mind. In the Price equation for class-structured populations, these three terms combine additively to give the evolutionary change in the mean phenotype. This article proposes a general formulation that clarifies this decomposition of the Price equation, both in discrete time and in continuous time. A key aspect of my treatment is that the evolutionary dynamics encapsulated by the Price equation are explicitly coupled with a set of equations describing the ecological dynamics through the dynamics of the vector **E**(*t*).

An influential idea in the theoretical literature, going back to Fisher (1930), is that the effect of selection is best captured by tracking the change in a weighted average rather than the more intuitive change in the arithmetic mean of the phenotype of interest. So far, this idea has been applied to exactly or approximately linear dynamics, where a focal population grows exponentially (Crow, 1979; Charlesworth, 1994; Engen et al., 2014). These systems are characterised asymptotically by a stable class structure (a right eigenvector of the constant projection matrix) and a stable set of reproductive values (a left eigenvector) associated with the long-term growth rate. Using these constant reproductive values as weights, the weighted density of the population grows *from the start* as it would when the stable class structure is reached. Furthermore, these constant weights can be used to cancel out the passive changes in the mean trait and therefore extract the signal of natural selection from the purely demographic consequences of class dynamics (Engen et al., 2014; Grafen, 2015b).

This article provides a general extension of this result, provided a dynamical and demographic definition of reproductive values is used. At a conceptual level, we need a clear distinction between types and classes, but to compute reproductive values we only need to work at the demographic level, using the between-class transition rates obtained by averaging over all types. The results hold for a large class of ecological models, allowing for density- and frequency-dependence, non-equilibrium population dynamics and environmental fluctuations. In addition, although I have focussed on discrete trait and state distributions, the derivation of Appendix A carries out unchanged if the trait averages are computed over a continuous distribution. This provides a direct connection with previous quantitative genetics models of age- and stage-structured populations (Lande, 1982a; Barfield et al., 2011). Furthermore, the result also extends to populations structured by continuous states, such as age-structured (Box 1) or size-structured populations studied by integral projection models (Rees & Ellner, 2016; results not shown). However, in practice, it may often be more useful to segregate a population into discrete classes, as this allows each class to be sufficiently populated.

The definition of reproductive values used in this paper departs from the classical usage in two ways. First, class reproductive values are not defined asymptotically, but as functions of time. However, the classical computation of reproductive values as an eigenvector of a constant projection matrix is obtained as a special case of the dynamical definition when the transition matrix for the ecological dynamics is constant. This occurs in particular when populations are at ecological equilibrium, as typically assumed in invasion analyses. Second, I emphasise a purely demographic notion of reproductive value. In particular, there is no need to assign a reproductive value to each genotype in the population. Rather, the relevant weights need to be calculated from the demographic dynamics where the genotype-specific vital rates are averaged within each class. This use of reproductive values contrasts with other definitions (e.g. Crow, 1979), but appears to match the definition attributed to Fisher (1930) by Grafen (2015a) and Grafen (2015b). Defining reproductive values at a demographic level allows one to circumvent the need for fitting models with phenotype- or genotype-dependent vital rates. Instead, we only need to estimate demographic projection matrices from the aggregated data where individuals of different genotypes are grouped by classes.

### 6.1 The different usages of reproductive values

An important question to ask is whether the properties of reproductive values discussed here are of relevance for practical studies of natural selection. The usefulness of reproductive value clearly depends on the biological question. First, one might be interested in detecting patterns of natural selection in demographic and genetic data, as collected for instance in field or controlled experimental studies. Then, it is possible to compute reproductive values by iterating estimated projection matrices backward in time, and use them as weights to detect deviation from neutrality. This use of reproductive values has been discussed by Engen et al. (2014), in the more restrictive setting of exponentially growing populations where reproductive values can simply be calculated as a constant eigenvector. In this article, I present an illustration of this approach using simulated data. Thus, if we are interested in understanding past events, reproductive-value weighting provides a useful way to test for the presence of selection without mistaking for selection the passive changes in mean trait resulting from class dynamics. Note that, although these passive changes are expected to disappear quickly in haploid linear models, in more realistic models, ecological feedbacks and genetic constraints may potentially sustain fluctuations in allele frequencies among classes on longer time scales, at least long enough for these fluctuations to become relevant for empirical or experimental studies. An example is given in figure 1, based on the classical LPA model for *Tribolium* dynamics. Haplodiploid systems of inheritance provide another example of this phenomenon (Gardner, 2015).

Alternatively, one may be interested in predicting patterns of evolutionary change for a particular trait of interest. If, for instance, one seeks to make predictions about how the virulence of a pathogen can be expected to change after the introduction of a vaccination campaign, the transient dynamics are of direct relevance to identify a potentially deadly short-term epidemic by a virulent strain that will eventually go extinct in the long run. Whether these changes are caused by natural selection or by class dynamics is a secondary issue. In addition, reproductive values can only be computed by backward iteration, so it is not clear how the concept is compatible with forward predictions on shortterm dynamics. For this type of forward-looking questions, the unweighted Price equation appears to be more useful. In particular, the unweighted Price equation arises naturally when studying short-term evolution in spatially structured population. For example, when studying the evolution of virulence during spatial epidemics on networks, Lion & Gandon (2016) found that the change in mean virulence depends on the build-up of a difference between the (local) virulence measured in hosts that have at least one susceptible neighbour and the (global) virulence measured at the population level. This term, which was interpreted as spatial differentiation in virulence, is the exact equivalent of the 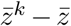 terms in equation (2).

For long-term evolution, the predictive power of reproductive values rests upon additional assumptions. For instance, if ecological dynamics take place on a fast time scale compared to evolutionary dynamics, the effect of transient ecological dynamics can be neglected and reproductive values can be computed on the ecological attractor. Thus, as for exponentially growing populations, we are interested in reproductive values in a “stable” population.
Equation (12) gives a general description of the dynamics of a weighted mean trait that can be combined with other genetic or ecological assumptions to derive expressions for the selection gradient. This suggests perspectives for analysing selection in polymorphic resident populations with arbitrary trait distributions. In addition, because most ecological models can be expected to be non-stationary (Chesson, 2017), a time-dependent concept of reproductive values may allow consideration of more complex, and realistic, population dynamics.

### 6.2 Neutrality, demography and selection

A key insight of equation (12) is that the effects of selection are captured by the covariance matrix **C** and weighted by the individual reproductive values, **v** and the class frequencies, **f**, which are purely demographic quantities computed using the average transition matrix **R**. In many problems in evolutionary game theory, the selection gradient is calculated by evaluating the change in mean phenotype due to a perturbation of a monomorphic population at equilibrium. The perturbation is caused by a new mutation, which is typically assumed to be rare, or to have a small effect on the phenotype (weak selection). Under these assumptions, the reproductive values and class frequencies in equation (12) can be approximated by those calculated in the resident monomorphic population, and this resident population, being monomorphic, may be thought of as “neutral”. However, real populations will often be polymorphic and characterised by a possibly multimodal trait distribution. It this case, it is still possible to ask how a mutation of small effect will affect the mean phenotype, but is is not immediately clear how the concept of a neutral reproductive value will be helpful. The results of this paper suggests that, for polymorphic populations with arbitrary trait distribution, the reproductive values and class frequencies in equation (12) should be approximated using the mean demographic matrix of the resident population. The reproductive values calculated in this way are still neutral because they are calculated in the resident population where the change in mean trait is assumed to be zero. However, they are not calculated under the assumption that the variance in the trait is vanishingly small. Instead, the standing genetic variation is handled by averaging over all types. Importantly, although the reproductive values do depend on the trait distribution of the resident population, they are not “selective reproductive values” because they are calculated from the average transition rates, so that information about the relative fitness of the different types is not used.

### 6.3 Possible generalisations

The derivation of the weighted Price equation also extends to multiple traits and environmental stochasticity. First, because the reproductive values do not depend on the trait one considers, the extension to several jointly evolving traits is straightfoward. However, potential correlations between traits will need to be accounted for in the transition rates. Second, the results extend directly to environmental stochasticity. In practice, if we have a random sequence of environmental variables **E**_*t*_ and associated demographic and genetic data, we can still use the backward recursion to compute reproductive values at different time steps, and then compute the reproductive-value-weighted mean forward. This is illustrated in figure 2. At a theoretical level, the asymptotic value of reproductive values under environmental stochasticity matches the results of Tuljapurkar (1989) in a density-independent model.

In contrast, the derivation is only valid for large populations of clonally reproducing types. More precisely, we need to have a sufficiently large number of individuals in each class. To account for the effect of small population sizes, we would need to model demographic stochasticity explicitly. Dynam-ical equations for reproductive values have been derived under demographic stochasticity (Rousset & Ronce, 2004; Lehmann, 2012), and this could provide a way forward. In principle, it should also be possible to extend the results to other genetic systems, including sexual reproduction or recombination, by using alleles as types and incorporating the genetic background into the class structure. Such potential extensions are left for future work.

Taking into account stochasticity is particularly important to fully extend the results of this paper to spatially structured populations. Many results on evolution in class-structured populations have been derived using an inclusive fitness formalism in subdivided populations (Taylor, 1990; Rousset, 2004; Lehmann & Rousset, 2014; Lehmann et al., 2016). In this approach, spatial structure is modelled using a deme-structured population and the local fluctuations in allele frequencies are taken into account through measures of population structure. It would be interesting to analyse how local stochasticity affects the definition and properties of reproductive values discussed here. However, if the total population size is sufficiently large, the spatial dynamics will follow approximately deterministic equations given by spatial moment equations (Lion, 2016; Lion & Gandon, 2016). The per-capita growth rates 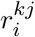 will then depend on the dynamics of higher-order spatial moments, and the equations for the mean trait and reproductive values will only represent the first in a hierarchy of dynamical equations. However, the key result of this paper would still be valid because it does not depend on any assumption on the functional form of the 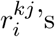.

Finally, I have assumed throughout that the trait under consideration can be measured in each class. This has clear limitations, because e.g. wing length cannot be used as a trait in the above formalism if we are studying an insect species with both alate and wingless state. This difficulty may be avoided by tracking the change in the frequencies of types, rather than the change in mean phenotype. Appendix A presents equations for the dynamics of the unweighted and weighted frequencies of type *i* in the population, with similar interpretations regarding the use of reproductive-value weighting. Working with frequencies should also be more appropriate to study a plastic trait that takes different values depending on the class in which it is expressed.

### 6.4 Connection with Fisher’s use of reproductive values

Historically, the use of reproductive values has also been advocated in two ways. In demography, reproductive values are often characterised as the weights *v*^*k*^ that need to be applied to the densities of each class (or age) so that the total reproductive value ∑_*k*_ *v*^*k*^*n*^*k*^ grows from the start with the longterm growth rate *r* (Fisher, 1930; Price & Smith, 1972; Samuelson, 1977; Crow, 1979; Charlesworth, 1994). The generality of this result has been debated, as this property of reproductive values seems tied to linear models (Samuelson, 1977; see also Bacaër & Abdurahman (2008) for an extension to periodic environments). However, the dynamical definition of reproductive values used here guarantees that the reproductive-valued weighted population size has the same growth rate as the non-weighted population size.

Alternatively, in evolutionary theory, reproductive values have been discussed in relation to Fisher’s Fundamental Theorem of Natural Selection (FTNS; Crow, 1979; Grafen, 2015a; Grafen, 2015b; Lessard & Soares, 2016), which states that the change in mean fitness *due to natural selection* is given by the genetic variance in fitness. In this literature, a focus of attention has been to determine whether Fisher’s intention in the FTNS was to use reproductive values as weights. In principle, we could obtain two different FTNS by substituting the growth rate *r*_*i*_ of type *i* for the trait *z*_*i*_ in the two Price equations derived above (Gandon & Day, 2009). However, these Price equations are derived for constant traits, whereas the growth rate *r*_*i*_ is a function of the environment **E**(*t*), and possibly of time itself if vital rates are functions of time. This will contribute an additional term to the Price equation, representing the feedback of the environment on the change in mean “fitness” (Frank & Slatkin, 1992; Gandon & Day, 2009; Lion, in press). Hence, as has long been recognised, the FTNS only captures a partial change in mean fitness, with or without reproductive-value weighting.

### 6.5 Summary

The results of this article confirm that reproductive values are best viewed as weights that can be used to decouple the changes due to selection from the passive changes due to demographic class dynamics. This allows one to measure selection in distinct classes with potentially different evolutionary values using a single, time-dependent currency. The practical interest of this approach is that the relevant weights at each time can always be calculated from time series, even for complex population dynamics.

## Acknowledgements

I thank Ophélie Ronce, Peter D. Taylor and five anonymous reviewers for very detailed and helpful comments on successive versions of this manuscript. This paper, matured during dark times, is dedicated to the memory of Joë l B.

## Appendix

### Appendix A: Derivation of the class-structured Price equation

#### A.1 No mutation

The mean trait in the *K*-class model is 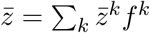 where *f* ^*k*^ = *n*^*k*^*/n* is the frequency of class *k*, and 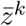 is the mean trait among individuals in class *k*. Introducing the frequency of *i*-individuals within class *k*, which is 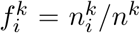, we have 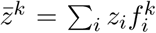. We first compute the dynamics of frequencies. Using the fact that 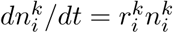, we have

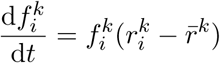

where the per-capita growth rate of type *i* in class *k* is

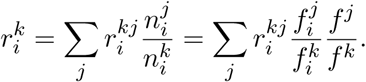

and the average per-capita growth rte of individuals in class *k*, averaged over all types, is

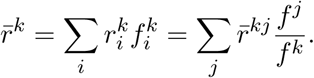

It is straightfoward to verify that 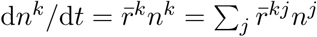 as expected.

Noting that 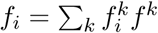, we have

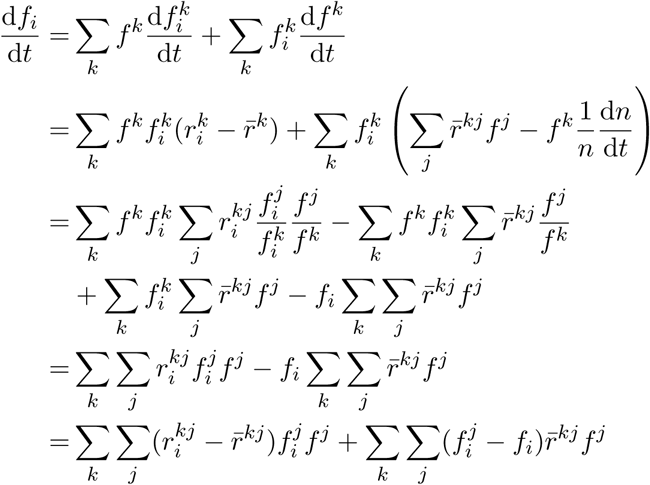

Multiplying by *z*_*i*_ and summing over *i* yields the dynamics of the mean trait

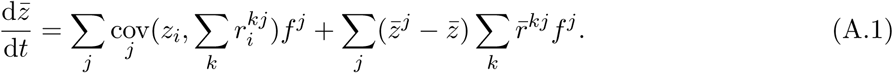

The dynamics of the mean trait in class *k* can be derived from the dynamics of 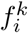. This gives

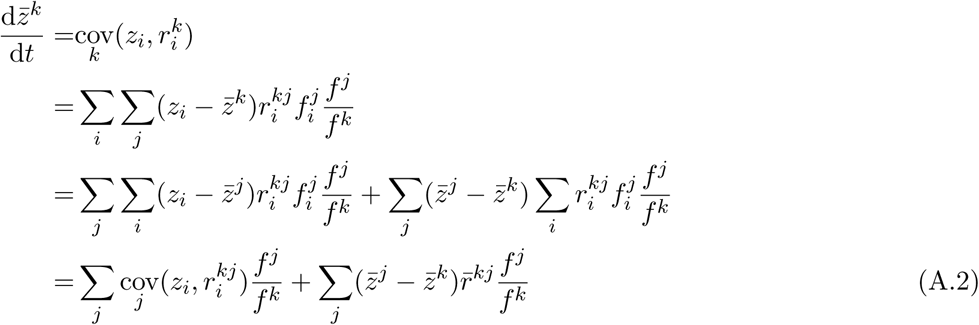

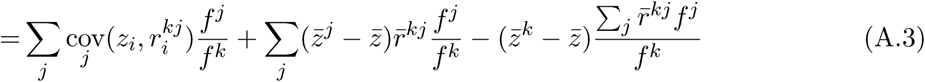

From equations (A.1) and (A.3), we can also derive the dynamics of 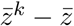 which gives:

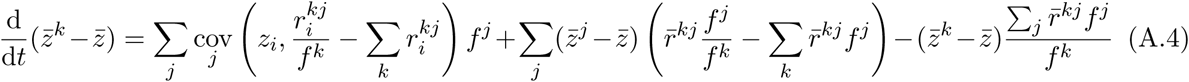

#### A.2 Mutation

Let us consider the following mutation model: mutations occur at rate *μ* and with probability *m*_*𝓁i*_ a parent of type *i* can give birth to an offspring of type *𝓁*, conditional on mutation. Assuming that the per-capita rate 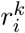 can be decoupled into birth and death contributions as 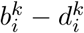, the change in the density 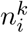 can then be written as

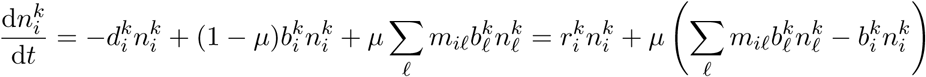

Thus, mutation contributes an additional term to the dynamics of 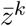

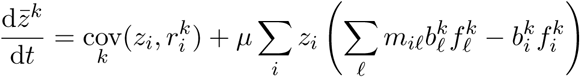

which can be rewritten as

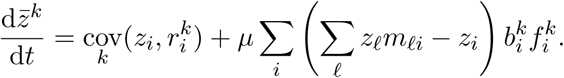

Hence, because 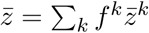, mutation contributes the following additional term to the dynamics of 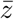

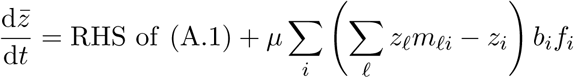

where 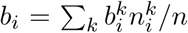 is the average birth rate of type *i* across all classes, and 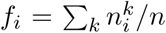 is the global frequency of type *i*.Note that the above derivation assumes that the rate and distribution of mutations are across classes.

Noting 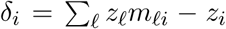 the deviation between parent and offspring trait, the second term of equation (A2) can be further split into two components, as follows:

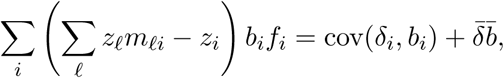

where 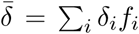 is the mean deviation over all types, 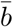 is the mean birth rate, and 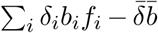 is the covariance between the trait difference and the birth rate. This is the continuous-time version of equation (12) in Barfield et al. (2011).

#### A.3 Weighted Price equation

We now calculate the dynamics of a weighted average frequency, 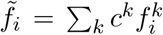, with weights *c*^*k*^(*t*) such that *c*^*k*^ = *v*^*k*^*f* ^*k*^ and ∑ *c*^*k*^ = 1. In the absence of mutation, this yields

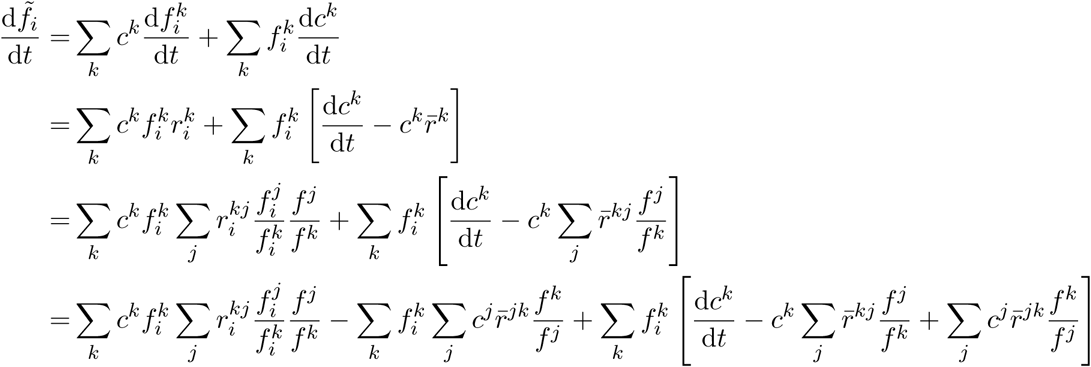

If the *c*^*k*^’s satisfy the system

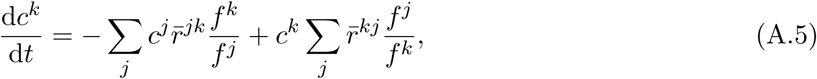

we then have the following simple equation for the dynamics of the weighted frequency

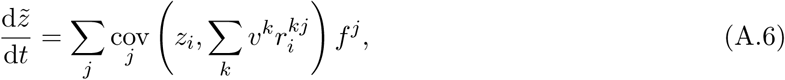

Multiplying by *z*_*i*_ and summing over *i* yields the dynamics of the weighted average 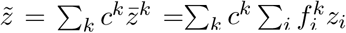

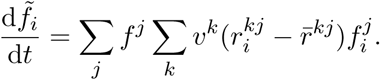

or in matrix form as

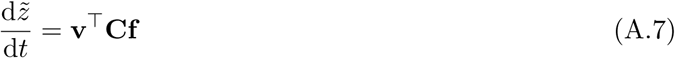

where **C** is the matrix of covariances with elements 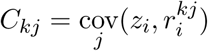. Taking into account mutation would only contribute an additional term, which is simply the second term of equation (A2) with 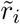 and 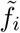 substituted for *r*_*i*_ and *f*_*i*_.

### Appendix B: Reproductive values

Equation (A5) can be rewritten in matrix form as

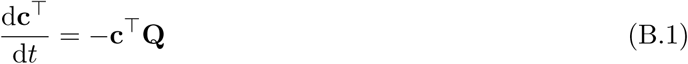

where the matrix **Q** has elements

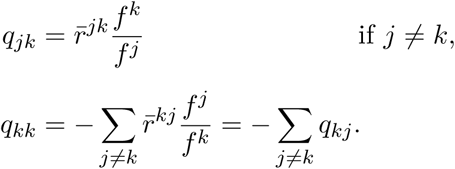

Similarly, we can find a dynamical equation for the *v*^*k*^’s. Because *c*^*k*^ = *v*^*k*^*f* ^*k*^ by definition, we have

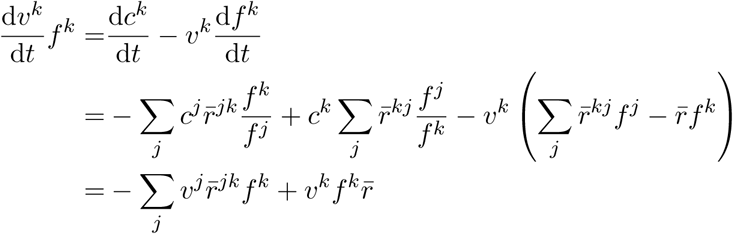

which gives us the following equation for the vector **v**

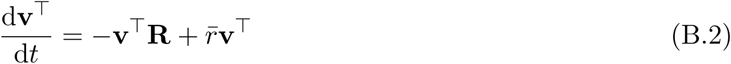

Equations (B.1) and (B.2) show that the vector **c** (resp. **v**) can be calculated at equilibrium as the left eigenvector of the matrix **Q** (resp. **R**), associated with eigenvalue 0.

### Appendix S: Supporting Information for “On the dynamics of reproductive values and phenotypic traits in class-structured populations”

#### S.1 Discrete time dynamics

Here I provide a derivation of the weighted and unweighted class-structured Price equations in discrete time.

##### S.1.1 Ecological dynamics

As for the continuous time, the ecological dynamics of a class-structured population are given by a matrix equation:

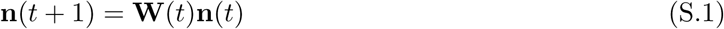

where **n**(*t*) is the vector of densities in each class, *n*^*k*^(*t*), and **W**(*t*) collects the quantities 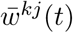. This gives us

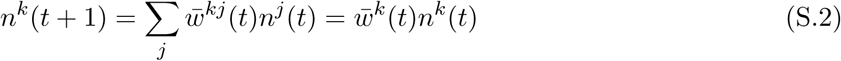

where 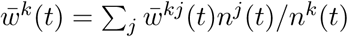. The total population size, *n*(*t*), obeys the following equation

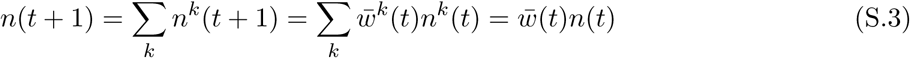

where 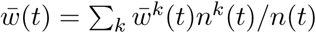.

Similarly, the dynamics of type *i* in class *k* can be written as

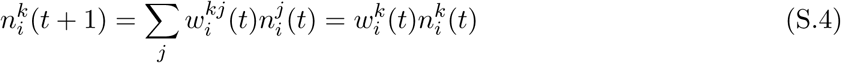

where

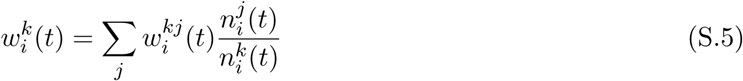

##### S.1.2 Change in frequency

The frequency of type *i* in class *k is 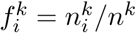*. The change in frequency is then

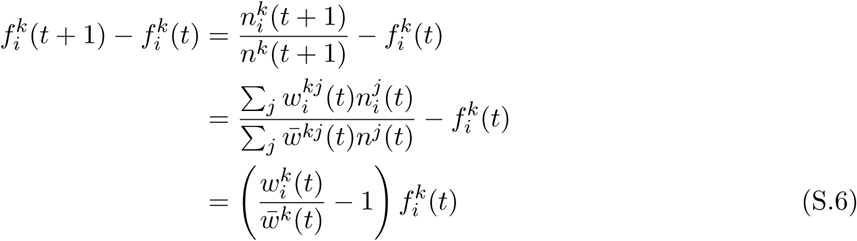

##### S.1.3 Change in mean trait

The change in the mean trait 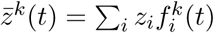 directly follows from the change in frequency:

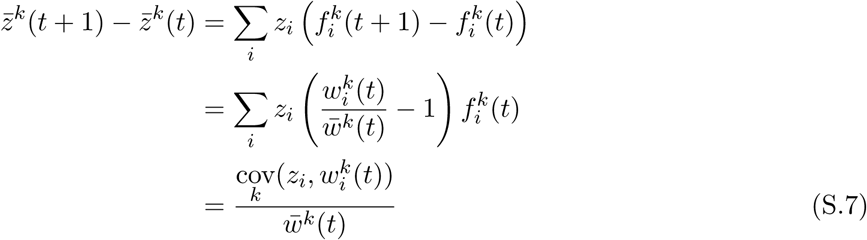

Using equation (S5), this can be expanded as follows

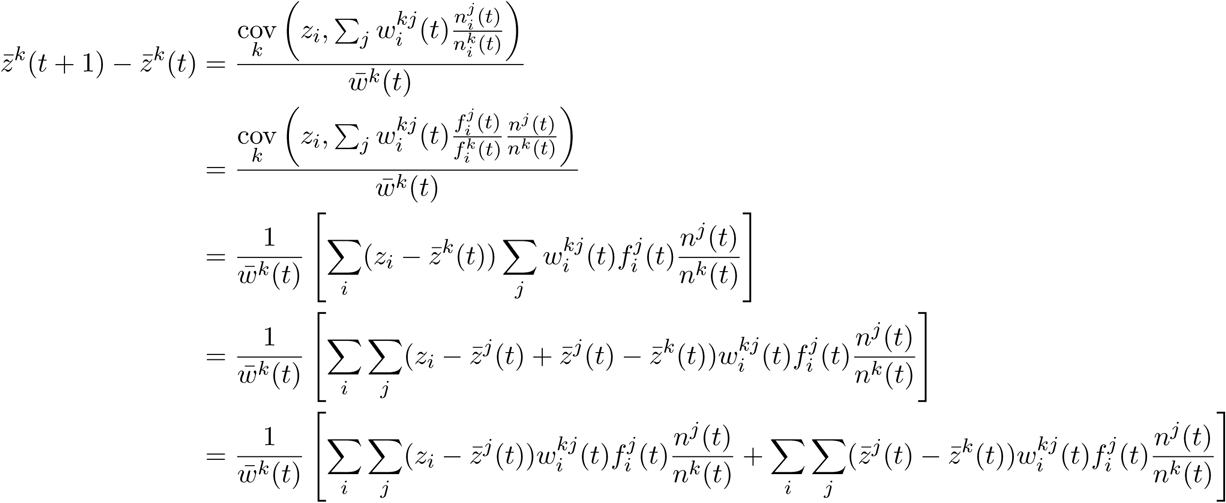

which gives finally

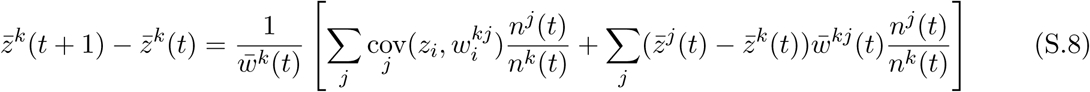

##### S.1.4 Change in weighted mean trait

We now introduce the following weighted average

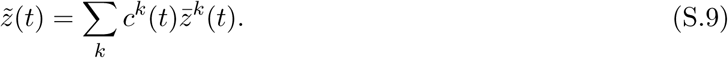

Using equation (S8), the weighted average at *t* + 1 can be written as

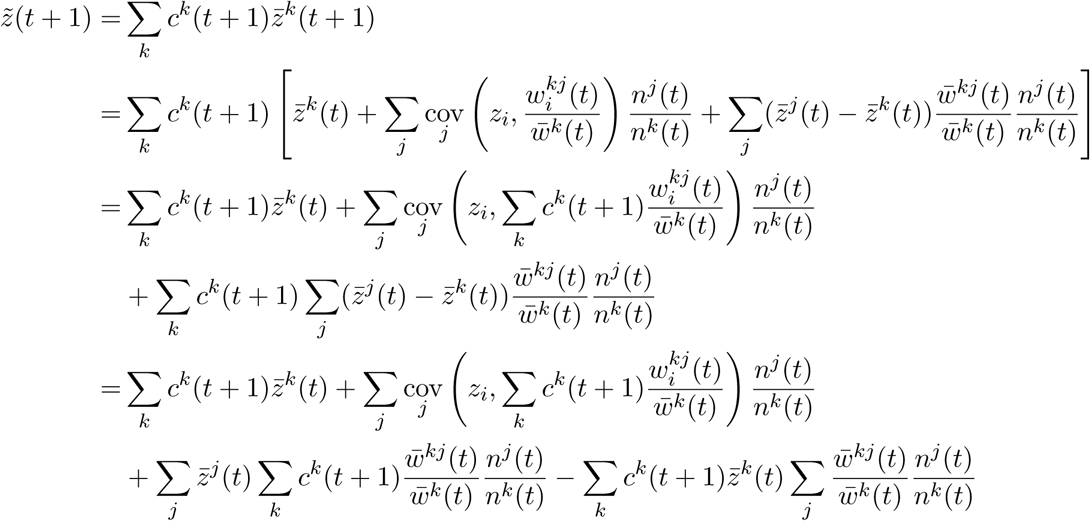

Because the sum over *j* in the fourth term is equal to one by definition, the first and fourth term cancel out and we obtain:

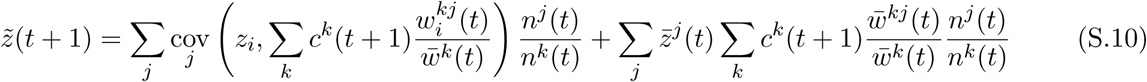

Now if we choose the weights *c*^*k*^ such that they satisfy the recursion:

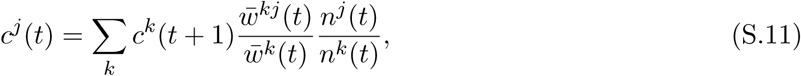

we obtain

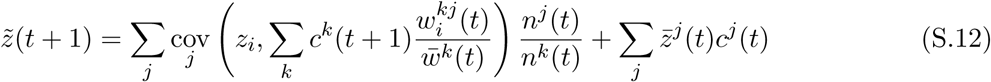

which gives us directly the change in the weighted average as

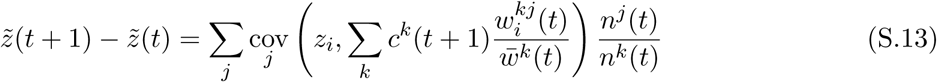

A final rearrangement uses the fact that 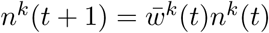 and the definition *c*^*k*^(*t*) = *v*^*k*^(*t*)*f* ^*k*^(*t*), so we have

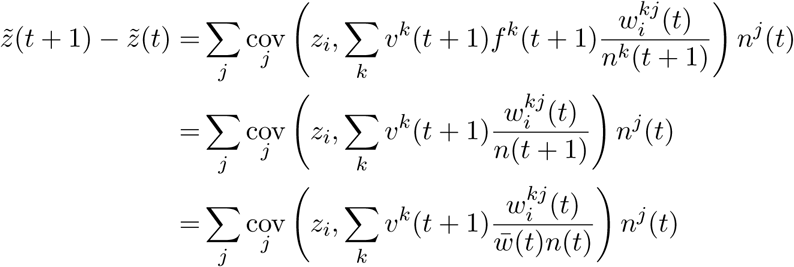

and we have finally

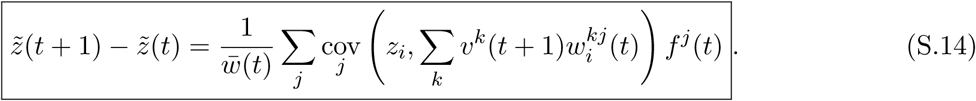

The latter equation thus shows that the change in the reproductive-value-weighted trait can be written as a covariance between the trait and a weighted measure of fitness, obtained by weighting each offspring in the next generation by the reproductive value of the class in the next generation.

A recursion for the individual reproductive can be obtained from equation (S11) by noting that 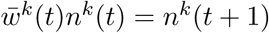. We then have, using the definition fo *c*^*j*^(*t*)

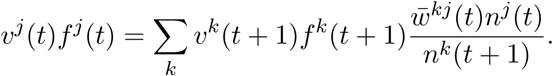

Because *n*^*k*^(*t*) = *f* ^*k*^(*t*)*n*(*t*), this can be simplified as

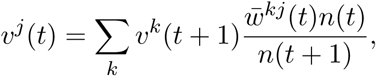

and using 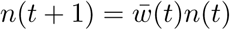 yields

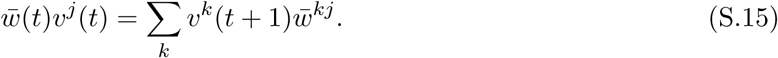

#### S.2 Derivation of the selection gradient for polymorphic and periodic resident populations

The starting point of the derivation is the equation for the dynamics of the weighted mean trait,

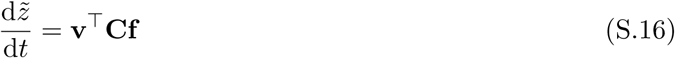

where the covariance matrix has elements

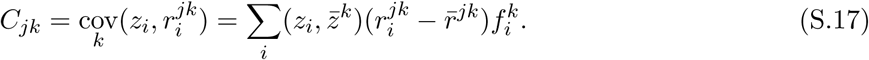

In a resident population at equilibrium, the change in the weighted mean trait is zero. We are interested in the perturbation resulting from the introduction of a mutation with small phenotypic effect.

##### Monomorphic population

In a monomomorphic population, the covariance matrix **C** is necessarily equal to the null matrix. Hence, in a population with two types, *w* and *m*, with the same trait value *z*_*w*_ the change in weighted mean trait is zero. If we now assume that the mutant type has trait value *z*_*w*_ + *ε*, we can write the resulting perturbation of the covariance matrix as **C** = **0** + *ε*^1^**C**^(^1^)^ + *ε*^2^**C**^(^2^)^ + *O*(*ε*^3^), where the **C**^(^1^)^ and **C**^(^2^)^ are the first and second-order perturbation terms of the matrix **C**.

If we assume that the resident population (when *ε* = 0) is on an ecological attractor, the perturbation of the reproductive values and class frequencies are 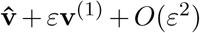 and 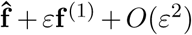, where 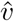 and 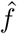 are calculated in the monomorphic population on its attractor. Plugging these expressions into equation (S16) yields the following perturbation of the change in weighted mean trait

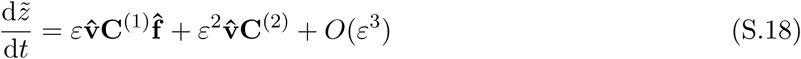

Hence, the leading-order term of the perturbation only depends on the perturbation of the covariance matrix, and not on the perturbation of the demographic variables **v** and **f**.

To calculate the perturbation of the covariance matrix, note that in a two-allele model, the relationship 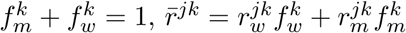 and 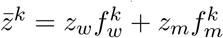 yield the following egality

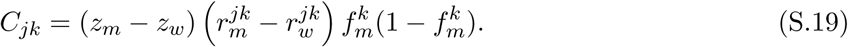

Now, if we assume that the effect of the mutation has only a weak effect on the ecological attractor, we can write the rates 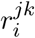 as a function of the trait *z*_*i*_ and of the resident monomorphic environment 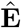. We then have

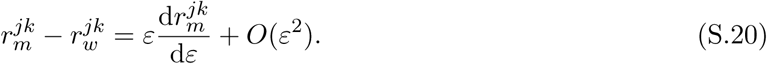

where the derivative is evaluated at *ε* = 0. Finally, this leads to

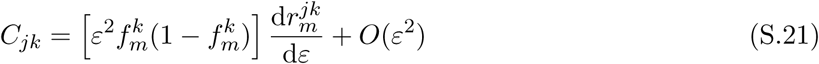

The factor between square brackets is the trait variance within class *k*, 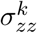. Under weak selection in a deterministic model, 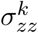 should be well approximated by the trait variance in the whole population, *σ*_*zz*_ (Lande (1982b); see also Barfield et al. (2011), their equation B11). Equation (S21) implies **C**^(^1^)^ = **0** and **C**^(^2^)^ = *f*_*m*_(1 *- f*_*m*_)d**R**_*m*_*/*d*ε*. This yields finally

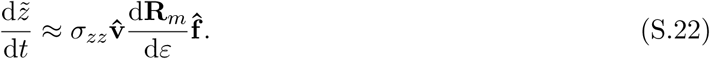

##### Periodic environment

The derivations of equations (S.18) and (S.21) make no assumptions on the nature of the resident attractor. Thus, equation (S22) can be applied to a periodic monomorphic attractor, but now the vectors 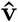 and 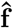 are time-dependent, and the matrix **R**_*m*_ depends on the timedependent resident attractor. In a deterministic model, the mutant frequency should not change in a neutral model, even in a periodic model, because the densities of the two types have exactly the same dynamics, so we can approximate the trait variance as a constant factor. Doing so, and integrating over one period gives the change in the weighted mean trait over one period (equation (23) in the main text).

##### Polymorphic resident populations

I now assume that *M* types coexist in the resident population at equilibrium. For such polymorphic resident populations, the perturbation of the covariance matrix can be calculated in different ways depending on how the mutation arises. For simplicity, I assume here that all individuals in type *m* mutate. The trait value in the mutant individuals is 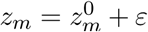. Assuming that the per-capita growth rates can be evaluated using the resident environment 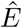, we write

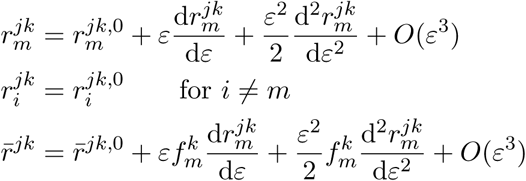

where the terms with a “0” superscript and the derivatives are calculated for *ε* = 0. In the following,we also note 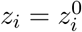 the trait value for *i ≠ m* and 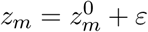 ; the mutant trait value. We then have 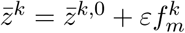 and

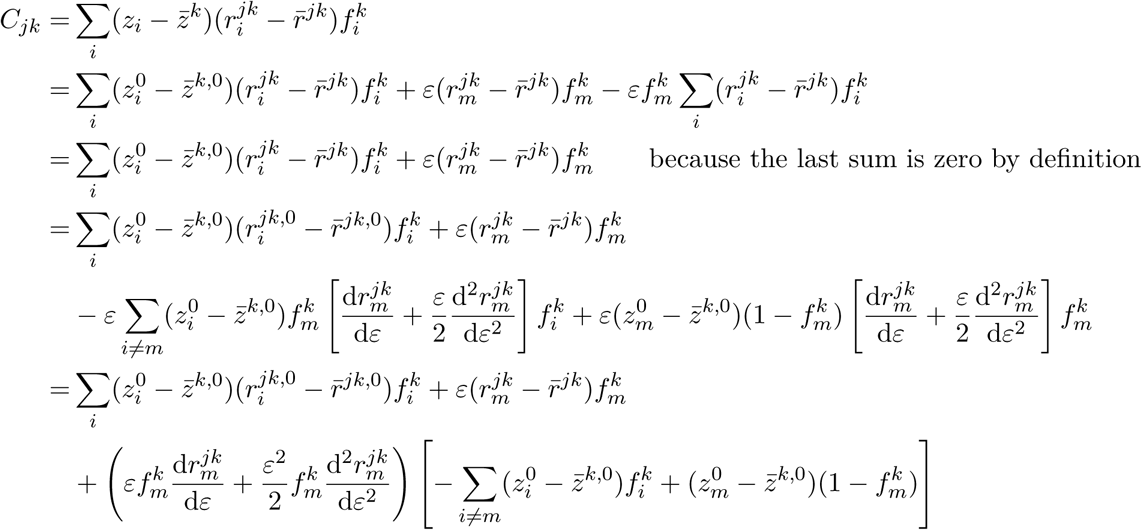

The first term is *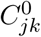*, the covariance evaluated at *ε* = 0. The term between brackets in the third term can be simplified as 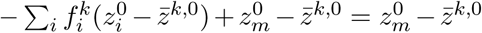. Expanding the second term then yields

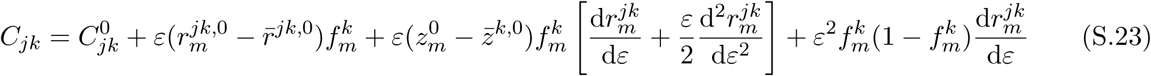

If the perturbations of the vectors **v** and **f** are assumed to be negligible compared to the perturbation of the covariance matrix, we can preand post-multiply by the vectors **v**^0^ and **f** ^0^ calculated at equilibrium in the resident population with *ε* = 0. We then have

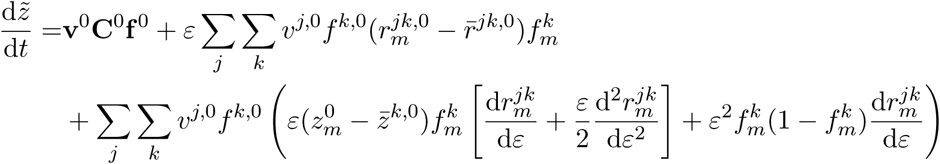

The first term is the change in mean trait in the resident population, which is zero because the resident population is at equilibrium. The second term can be simplified by noting that

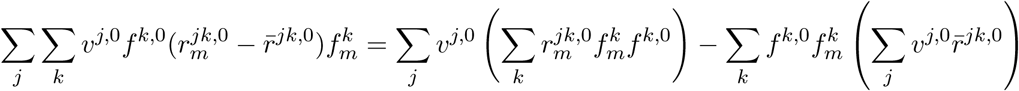

The first term between brackets is zero because it is proportional to 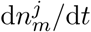 calculated for a neutral mutant. The second term between brackets is zero by definition of **v** in the resident population at equilibrium. Hence, the dynamics of the weighted mean trait can be approximated as

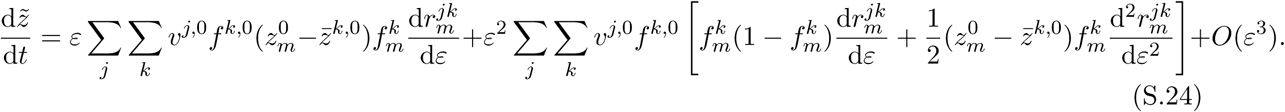

Importantly, the leading order term is *O*(*ε*), which is of the same order as the perturbations of **v** and **f**. Hence, to neglect the perturbation of demography compared to the perturbation of the covariance matrix, we need to make a more stringent assumption than in the monomorphic case. If this assumption does not hold, we need to compute the perturbation of the reproductive values and class frequencies.

Note that if the population consists of only two types *w* and *m* with traits *z*_*w*_ and *z*_*m*_ = *z*_*w*_ + *ε* (so that, when *ε* = 0, the population is monomorphic), equation (S22) can be recovered using the relationship 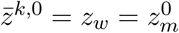.

#### S.3 Continuous age structure and Fisher’s original concept of reproductive value

Consider a population with continuous age structure. The density of type-*i* individuals with age *a* at time *t* is *n*_*i*_(*a, t*). These individuals die at rate *d*_*i*_(*a, t*) and give birth at rate *b*_*i*_(*a, t*). These assumptions yield the following partial differential equation

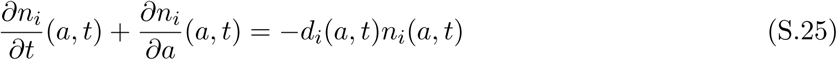

along with the boundary condition

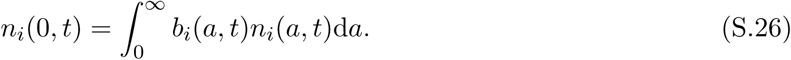

The total density of individuals at age *a* and time *t* is *n*(*a, t*) = ∑_*i*_ *n*_*i*_(*a, t*).

Now, for a focal trait *z* with value *z*_*i*_ in type-*i* individuals, we note 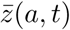 the average trait value in age-*a* individuals at time *t*. Noting *c*(*a, t*) the reproductive value at age *a* and time *t*, we calculate the weighted average

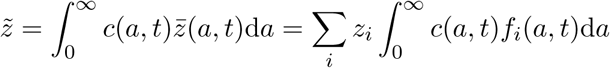

where *f*_*i*_(*a, t*) = *n*_*i*_(*a, t*)*/n*(*a, t*). In this specific case, it is easier to work with the individual reproductive values. Using the relationship *c*(*a, t*) = *v*(*a, t*)*f* (*a, t*) where *f* (*a, t*) = *n*(*a, t*)*/n*(*t*) is the frequency of individuals with age *a* at *t* in the population, we write

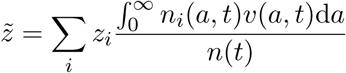

With these assumptions, the dynamics of the weighted mean trait is

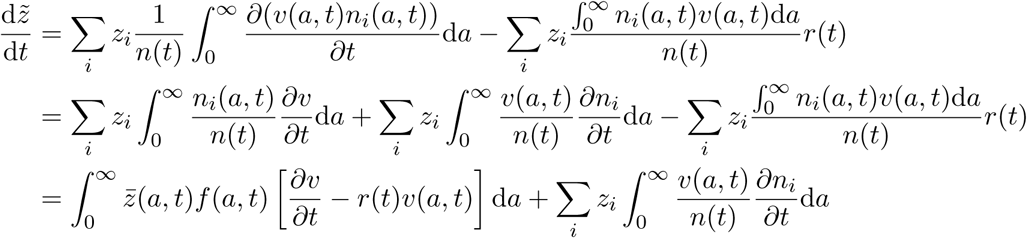

where *r*(*t*) = d ln(*n*)*/*d*t* is the per-capita growth rate of the total population.

Now using equation (S25), the last term becomes

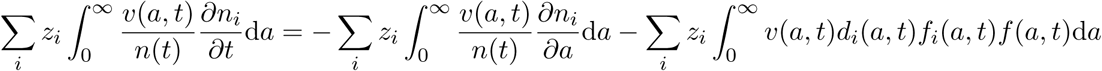

The first term on the right-hand side of the latter equation can be integrated by parts, which yields

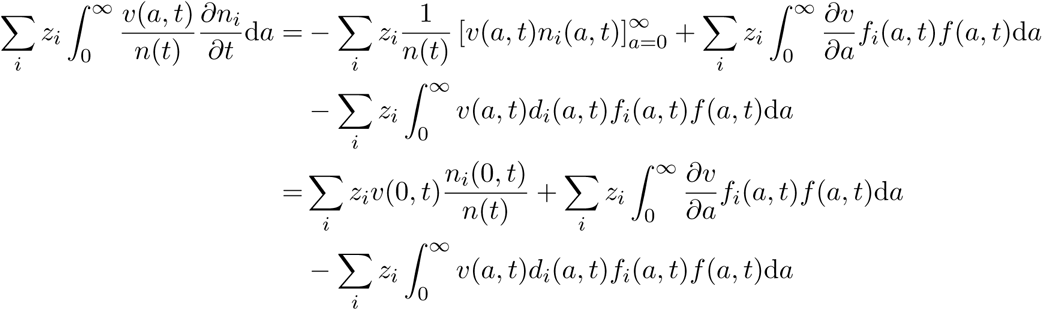

Plugging this into the dynamics for 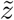 gives

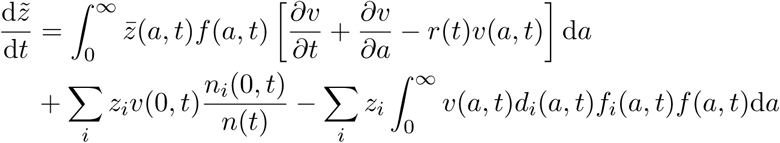

Using the boundary condition (S.26) for *n*_*i*_(0*, t*), we obtain

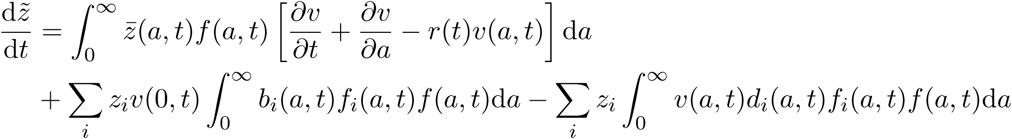

Finally, we introduce the age-specific covariances:

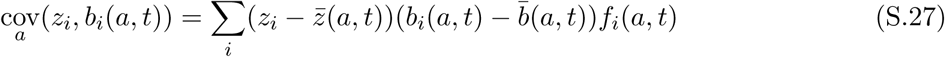

and similar expressions for the covariances between the trait and the death rate. Doing so results in the following equation

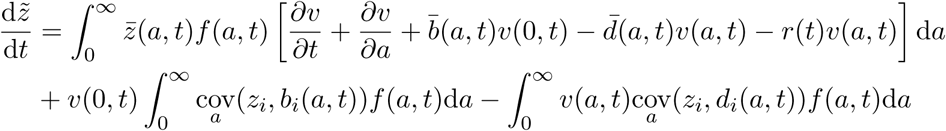

Thus, if the individual reproductive values satisfy the following partial differential equation

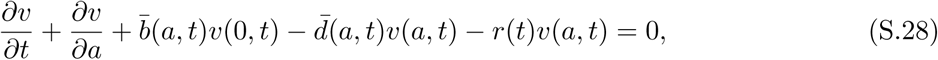

the dynamics of the weighted mean trait take the simple form:

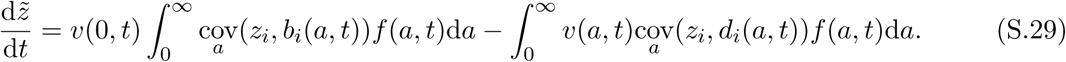

This is the equivalent of equation (S16) for a continuous-age structure. Note that the selective effects due to birth events are weighted by the reproductive values of newborns, *v*(0*, t*) and the frequency of adults with age *a*, *f* (*a, t*), whereas the selective effect due to death events at age *a* are weighted by the reproductive value at age *a*, *v*(*a, t*), and the frequency *f* (*a, t*).

Equation (S28) was previously derived by Bacaër & Abdurahman (2008) in a periodic epidemiological model structured by infectious age (their equation (7)). Equation (S28) generalises their finding to polymorphic population. As in Bacaër & Abdurahman (2008), equation (S28) is coupled to an adjoint partial diffrential equation for *f* (*a, t*), with normalisation condition 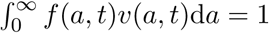.

Note that it is straightforward to show that, if the reproductive values satisfy equation (S28), the weighted population size 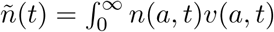 d*a* always grows as

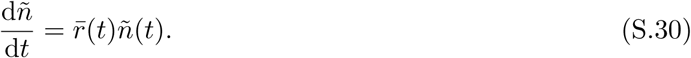

A special solution of equation (S28) can be found under the assumption that the birth and death rates are independent of time. Then, the solution of equation (S28) is

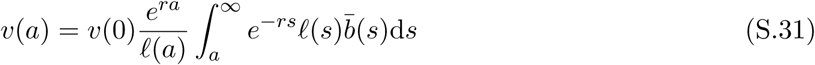

With 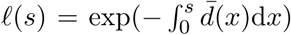 the probability to survive to age s.^2^ Expression (S.31) is the original definition of reproductive value given by Fisher (Fisher, 1930; Charlesworth, 1994), but with average birth and death rates. Note that in Fisher (1930)’s first version, the normalisation constant v(0) was omitted but this was corrected in the 1958 version of the book.

## Footnotes

1 More generally, the matrix R can depend on a density-independent ergodic environment (Tuljapurkar, 1989).

2 An equivalent expression, with time-dependent coefficients, can be derived if the age distribution is assumed to stabilise quickly relative to the time scale on which 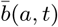 and 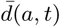 change, see e.g. Day et al. (2011).

